# Camelina CircRNA Landscape: Implications for Gene Regulation and Fatty Acid Metabolism

**DOI:** 10.1101/2024.07.02.601705

**Authors:** Delecia Utley, Brianne Edwards, Asa Budnick, Erich Grotewold, Heike Sederoff

## Abstract

Circular RNAs (circRNAs) are closed-loop RNAs forming a covalent bond between their 3’ and 5’ ends, the backsplice junction (BSJ), rendering them resistant to exonucleases and thus more stable compared to linear RNAs. Identification of circRNAs and distinction from its cognate linear RNA is only possible by sequencing the BSJ that is unique to the circRNA. CircRNAs are involved in regulation of their cognate RNAs by increasing transcription rates, RNA stability and alternative splicing. We have identified circRNAs from *Camelina sativa* that are associated with the regulation of germination, light response, and lipid metabolism. We sequenced light-grown and etiolated seedlings after 5 or 7 days post-germination and identified a total of 3,447 circRNAs from 2,763 genes. Most circRNAs originate from a single homeolog of the three subgenomes from allohexaploid camelina and correlates with higher ratios of alternative splicing of their cognate genes. A network analysis shows the interactions of select miRNA:circRNA:mRNAs for regulation of transcript stabilities where circRNA can act as a competing endogenous RNA. Several key lipid metabolism genes can generate circRNA and we confirmed the presence of KASII circRNA as a true circRNA. CircRNA in camelina can be a novel target for breeding and engineering efforts.

**Core ideas:**

1. First discovery of 3,447 genic and 307 intergenic unique putative circRNAs from *Camelina sativa*.
2. We identified circRNAs that were regulated in response to seedling de-etiolation.
3. Most circRNAs originate from only one homeolog of the three subgenomes in this allohexaploid Camelina.
4. Alternative splicing of exon skipping and intron retention positively correlate with circRNA occurrence.
5. Validation of KASII circRNAs as an example of lipid metabolism pathways potentially regulated by circRNA.

## 1. INTRODUCTION

In recent years, endogenous circular RNAs (circRNAs) have been identified as regulators of transcription and translation in eukaryotic cells. Circular RNAs are thought to be formed during co-transcriptional pre-mRNA processing in the spliceosome, where a downstream 5’ splice site is covalently linked to an upstream 3’ splice site generating the backsplice junction (BSJ) as reviewed in (C.-X. Liu & Chen, 2022). The generation of circRNAs from their cognate linear transcripts is cell-type specific and is controlled by developmental and environmental changes. While in animal cells sequence elements, like inverted repeats, have been identified in some cognate genes to be involved in circularization, those elements are less common in plants (Ye et al., 2015). The pathways and regulation by which circRNA biosynthesis is generated are still unknown as are the enzymes involved in the circularization. The molecular functions of circRNAs are diverse and involve the regulation of alternative splicing, serving as scavengers for miRNAs and RNA binding proteins, and the regulation of transcription and translation of their cognate genes. In animal cells, some circRNAs are translated into short peptides that can disrupt signaling by mimicry of protein:protein binding sites (C.-X. Liu & Chen, 2022).

In plants, only a few circRNAs have been functionally characterized and shown to regulate transcript abundance via miRNA sponging (Y. Liang et al., 2019), and alternative splicing via R-loop formation (Conn et al., 2017). Despite their potentially important roles in gene expression and possible use as targets for the genetic improvement of crop production, circRNAs have only been sequenced from about 20 plant species including Arabidopsis, rice, and bamboo. This lack of circRNA identification is not only due to its novel discovery but also its very low abundance and high potential for artifacts (Budnick et al., 2024; Nielsen et al., 2022). The circRNA molecule is only distinguishable from its linear cognate RNA by the BSJ sequence. Different methods have been developed to identify circRNA via the BSJ sequence. These methods utilize various combinations of circRNA enrichment from total RNA pools by specific digestion or pull-down of linear RNA, followed by amplification and either long-read (Nanopore) or short-read (Illumina) deep sequencing (Budnick et al., 2024; Z. Liu et al., 2021; Rahimi et al., 2021). Because different genetic events and features as well as technological inaccuracies can lead to the identification of relatively large amounts of false BSJ reads and therefore false positive circRNAs (Nielsen et al., 2022), it is essential to validate circRNAs via RT-PCR using divergent and convergent primer sets on cDNA and gDNA samples from the tissue (Budnick et al., 2024; Dodbele et al., 2021).

CircRNAs have been sequenced from over 20 plant species to date, but not from any oil seed crop (Chu et al., 2017). The closest genetic relative to Camelina spp. with publicly available circRNA sequence information is *Arabidopsis thaliana,* a diploid reference model plant from the Brassicaceae family (Philips et al., 2020). CircRNAs have also been reported from different tissues of the diploid *Brassica rapa* (Chinese cabbage), identifying a total of about 1,300 circRNAs (Wang et al., 2019). No oilseed-producing Brassica species has been sequenced for circRNA detection.

A high-quality reference genome is available for the allohexaploid Camelina genome (Kagale et al., 2014). Camelina is closely related to the diploid *Arabidopsis thaliana* and more distantly to the oil seed crop canola. Its polyploidy originated from a triplication event, as a comparison to Arabidopsis showed (Kagale et al., 2014), resulting in three nearly identical copies of genes that are orthologous to many single-copy genes in Arabidopsis Transcriptome analysis showed that 77% of all genes were retained as triplicated homeologs, with an expression level advantage of one of the subgenomes (Cs-G3). Over 80% of the 772 non-redundant genes encoding proteins involved in fatty acid and lipid metabolism (FALM) in Camelina were retained in triplicate. A subset of those FALM genes experienced further expansion, with some exceeding >10 paralogs.

Fatty acid and lipid metabolism in plants has been intensely studied because products like seed oil are a major source of human nutrition and have many industrial applications, but also because they play essential roles in plant growth, development, signaling, and stress responses in addition to their structural importance as membrane components (Y. Liang et al., 2023; Li-Beisson et al., 2013). The enzymes of these pathways are regulated at the transcriptional, translational, and posttranslational levels. Fatty acid and lipid synthesis, storage, modifications, and degradation underlie coordinated regulation between all cellular compartments and cell types (Li-Beisson et al., 2013; Schmid, 2021). Many fatty acids also serve as substrates for waxes, hormones, and modifiers of proteins or other polymers (Kumar et al., 2022; Lewandowska et al., 2020; Weber, 2002).

Camelina has already been recognized as an excellent source of drop-in replacement fuel due to its high seed oil content and its ability to grow with low input on marginal lands, especially under drought conditions. Its utility for second-generation fuels has been demonstrated, and Camelina meal is approved and used as a feed supplement, providing additional value and reducing the fuel-versus-food competition (Neupane et al., 2022). Engineering and breeding efforts have successfully generated improved varieties and transgenic/gene-edited lines (Dalal et al., 2015; Edwards et al., 2022; Kawall, 2021, 2021; Neumann et al., 2021; Wilson et al., 2022). Despite all these improvements, commercialization is still lagging due to variability in oil production in response to environmental factors.

We are presenting here a transcriptome analysis of camelina seedlings, distinguishing linear and circular RNAs. To identify a wide set of circRNAs involved in many aspects of development and general lipid metabolism, we chose to sequence young seedlings that were either etiolated after 5 or 7 days in the dark or de-etiolated and photosynthetically active after diurnal exposure to light. These two contrasting treatments and time points have distinct fatty acid and lipid synthesis and degradation profiles as etiolated seedlings utilize seed oil for survival and development while the light-grown photosynthetically active seedlings mobilize the storage oil but also synthesize their own fatty acids de-novo (Li-Beisson et al., 2013). We identified linear and circular RNA from rRNA-depleted cDNA that was generated with random primers and sequenced using Illumina short-read technology (Budnick et al., 2024). These data were used to identify light-regulated gene expression for linear and circular transcripts. We identified 3,447 circRNAs from camelina seedlings. Qualitatively and quantitatively there were more circRNAs in the etiolated seedlings compared to the light-grown ones. Comparison of homeologs showed that 62% of all circRNAs originated from only one homeolog, but some circRNAs were generated from two or even all 3 homeologs. Some highly abundant circRNAs were strictly conserved among all samples, while a smaller number of circRNAs were identified as treatment-specific. Correlation with the linear transcripts showed that a higher percentage of circRNAs originated from transcripts with alternative splice (AS) variations in intron-retention or exon-skipping. Some circRNAs contained putative miRNA binding sites and could serve as competing endogenous RNAs (ceRNAs) to stabilize linear RNAs. Lipid metabolism genes were identified as circRNA generating genes and a comparison to Arabidopsis, soybean and rice showed that some of those identified are conserved. As an example, we confirmed the circRNAs identified from all 3 Camelina KASII homeologs as true circRNAs. The identified Camelina circRNAs can serve to identify new regulatory mechanisms in fatty acid and lipid metabolism in oil seed crops.

## 2. MATERIALS AND METHODS

### 2.1. Plant Material

Wildtype *Camelina sativa* (cultivar Calena) seeds were sterilized in a solution containing 0.5× bleach and 50μl of Tween-20 for 10 minutes and rinsed with sterile distilled water. Approximately 45 sterilized seeds were placed onto 0.5× Murashige and Skoog media plates containing, 0.8% agar. Plates were divided equally into four separate cardboard boxes wrapped in aluminum foil to prevent light contamination. Wrapped boxes received 24 hours of dark stratification at 4°C before being moved into treatment conditions. Following dark stratification treatment, plates were moved into a temperature-controlled growth chamber with a 21℃/18℃ light/dark temperature cycle and 16-hr day length. For the light treatment, plates were removed from the foil-wrapped box and distributed onto a growth chamber rack for 5 or 7 days before harvesting. Dark-treatment plates remained in a foil-wrapped box for the remainder of the experiment until harvest. For tissue collection, seedlings from both treatments were removed from the growth chamber at the same time of day and dark-treated plates were unwrapped and harvested in the dark using a green LED headlamp. Seedlings from the same petri plate were treated as one sample.

### 2.2. RNA extraction and Sequencing

Total RNA was extracted from pulverized seedling tissue using the PureLink RNA mini kit (ThermoFisher Scientific) following the manufacturer’s instructions and contaminating gDNA was digested using the TURBO DNase-free kit (ThermoFisher Scientific). RNA quality was evaluated using a BioAnalyzer 2100 Plant RNA Nano assay (Agilent) and samples with RIN > 8.5 were selected for downstream library prep and sequencing. Samples were rRNA depleted using the QIAseq FastSelect –rRNA Plant Kit (Qiagen) before library preparation using the NEBNext Ultra II Directional RNA Library Prep Kit for Illumina (New England Biolabs) and sequencing on the NovaSeq6000 (Illumina).

### 2.3. Transcriptome analysis and identification of circRNA

As circRNAs are known to be associated with alternative splicing events (Conn et al., 2017), the discovery of novel transcripts and unannotated isoforms was of particular interest. Thus, quantification of all other (non-circular) RNA transcripts was performed using an assembly tool that enables quantification of mapped fragments regardless of whether or not they follow the form defined by the standard genome annotation file. Fragments that aligned to the Camelina reference genome in the first step of the CLEAR pipeline and resulted in distinct, primary alignments were used as input for transcript assembly using StringTie (v. 2.2.1) (M. Pertea et al., 2016). Initial transcript assembly was performed for each sample separately based on HISAT2 read alignments using Camelina genome annotations (https://cruciferseq.ca/Csativa_download, version 2.0) as a guide. Assemblies for all samples were merged into a single assembly containing all known transcripts, novel transcripts, and transcripts with retained introns. Transcript abundances were then re-estimated for each sample using the final, merged assembly as the reference genome as a guide. Attributes of novel transcripts were compared to the known, annotated transcripts using GFFcompare in the GFF Utilities package (G. Pertea & Pertea, 2020). Differential expression analysis was performed at the transcript level with fragments per kilobase of transcript per million reads sequenced (FPKM) as the expression measurement using the Ballgown statistical analysis package in R (Frazee et al., 2015).

To identify total transcript abundances from differentially expressed genes (DEGs), files generated by StringTie (M. Pertea et al., 2015) (v2.2.1), were used as input for prepDE.py (provided in StringTie manual) to generate a matrix of read counts. The counts matrix was used as input for edgeR (Robinson et al., 2010). Genes with ≤ 10 reads were discarded, and counts were normalized by effective library size. Transcripts that were not assigned to a known Camelina gene ID were discarded. P-values were calculated on normalized counts and a False Discovery Rate (FDR) threshold was applied to adjust for multiple comparisons.

Identification of circRNA was completed using two independent detection pipelines: CIRI2 (Gao et al., 2015, 2018) and CLEAR (Ma et al., 2019a), as described in previous work (Budnick et al., 2024; J. Zhang et al., 2020).

PANTHER classification (v18) (Mi et al., 2019) was used for identification of Biological Process GO terms enriched among circRNA parent genes. GO annotations from homologous Arabidopsis genes were used as input. Using the initial PANTHER output, top parent categories with significant overrepresentation and FDR < 0.05 were selected and visualized in R.

### 2.4. PCR confirmation and Sanger sequencing

CircRNAs and their cognate transcripts from lipid metabolism genes were validated using RT-PCR with convergent and divergent primers (Supplementary Table S1). DNase-treated total RNA seedlings were used for first strand cDNA synthesis, performed using SuperScript IV Reverse Transcriptase with random hexamers (Thermofisher Scientific) following the manufacturer’s protocol. Divergent primers were designed to generate PCR products spanning the BSJ and to capture as much of the internal sequence of the circRNA as possible. Primers for CsActin2 (Csa15g026420) were used as a linear positive control. RNAse-A treated genomic DNA (gDNA) was used for false positive controls to identify genomic rearrangements. PCR on both cDNA and gDNA were amplified using OneTaq Hot Start 2X Master Mix (New England Biolabs). cDNA reactions were done using 10% total volume of the reaction as input and 100 ng of gDNA as input. Cycling conditions followed the manufacturer’s protocol. CircRNAs were allowed an extension time of 20 seconds while for longer circRNAs, controls, and linear products had an extension time of 30 seconds. PCR products were visualized by agarose gel electrophoresis. The PCR products from divergent primers of expected length were excised and purified via PCR clean up kit (New England Biolabs) for confirmation of BSJ using Sanger Sequencing.

### 2.5. Prediction of miRNA Targets

Sequences of mature microRNA from *Camelina sativa* were downloaded from the miRBase public database (Griffiths-Jones et al., 2006; Kozomara & Griffiths-Jones, 2014) and used to identify circRNAs with putative miRNA binding sites. CircRNAs expressed from genes that Poudel et al. identified as predicted targets for miRNAs in *Camelina sativa* were selected for miRNA target prediction (Poudel et al., 2015). Genomic sequence between the BSJ of each of these circRNAs was extracted and psRNATargetV2 was used to predict miRNA targeting with default parameters (Dai et al., 2018). psRNATargetV2 scores the complementation of a small RNA and potential targets and also considers the unpaired energy of the target region to predict potential binding sites. The full set of *C. sativa* coding sequences was also analyzed for putative miRNA targets in the same manner in order to build potential competing endogenous RNA networks of circRNA-miRNA-mRNA interactions. A subset of putative miRNA-target circRNAs came from genes which were implicated in cytoskeletal systems and a network was made using a custom R script to visualize the interconnections between circRNA and CDS targets in the network.

## 3. RESULTS

### 3.1. Phenotypic effects of light/dark germination of camelina seedlings

Seedlings germinated under constant darkness (Dark) or 16h:8h light/dark cycles (Light) for 5 or 7 days after stratification showed the characteristic phenotypic differences of etiolated and light-grown plants (Figure 1). The seedlings were flash frozen in liquid nitrogen prior to RNA extraction and sequencing library preparation.

**Figure 1:**
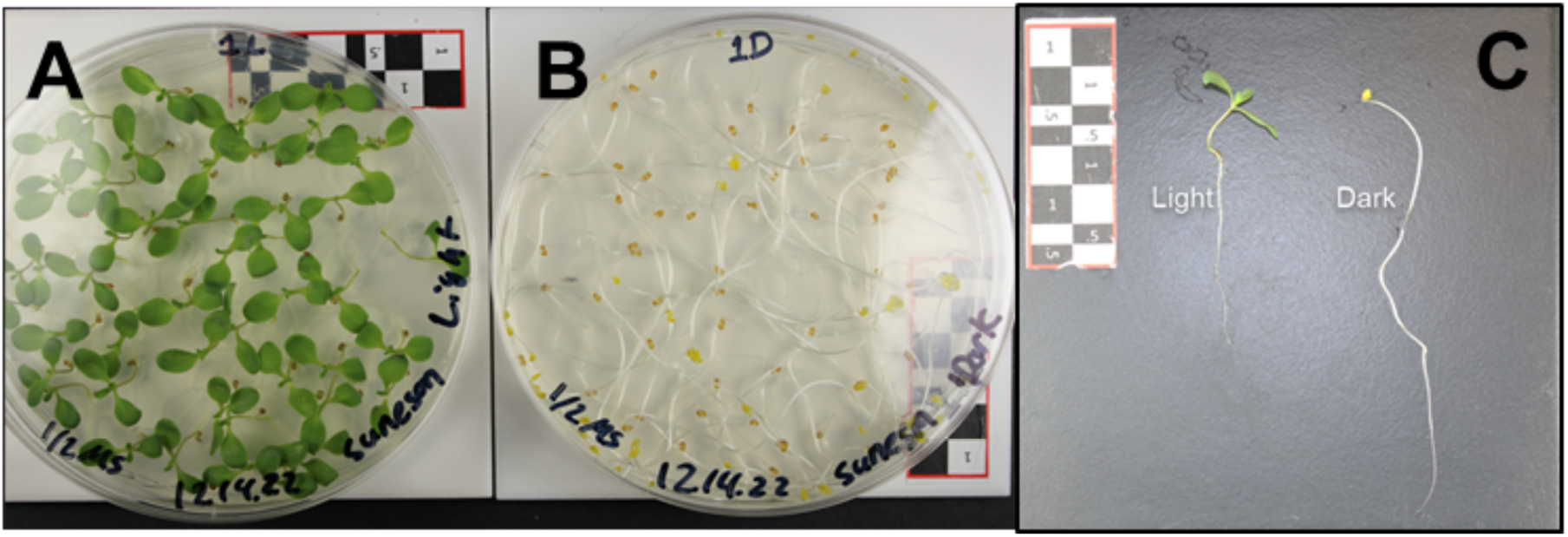
*Camelina sativa* seedling phenotypes after exposure to 16:8hrs light:dark cycles (A; Light) or continuous darkness (B; Dark) for 7 days post-stratification. (C) Etiolated seedlings show the “classic” elongated hypocotyl, hook, lack of chlorophyll, and long roots, while light-exposed seedlings show the emergence of the first pair of true leaves and shorter roots.

### 3.2. Sequencing and identification of linear and circular RNA

Circular RNAs are known to be at low abundance compared to linear RNA, which requires deep sequencing to identify sufficient numbers of reads containing BSJs, the only sequences that distinguish circRNAs from their linear cognate transcripts. Total RNA was extracted from four bioreps, each consisting of about 45 seedlings grown for five- and seven days under light:dark (16h:8h) or in continuous darkness (Figure 1). Ribosomal RNA-depleted sequencing libraries were generated using random hexamer primers and sequenced on Illumina NovaSeq6000 with a target of 60 million reads per sample. 150 bp paired-end reads were assessed for quality using FASTQC and only high-quality reads (Q>30) were used in downstream analyses. Overall sequencing depths ranged from 43 to 87M reads across all samples with an average of 50M and above per treatment group (Supplementary Table S2). Reads were aligned to the Camelina genome (https://cruciferseq.ca/Csativa_download) using the splice-aware aligner HISAT2 (D. Kim et al., 2019) with recommended alignment parameters for simultaneous identification of circRNA and alignment of linear RNA using the **C**ircular and **L**inear RNA **E**xpression **A**nalysis from **R**ibosomal RNA (CLEAR) pipeline (Ma et al., 2019). Following the standard guidelines for the bioinformatic discovery of circRNA, a separate orthogonal method of circRNA identification was applied using the CIRI2 pipeline (v2.0.6) (Gao et al., 2015). Untrimmed read pairs were aligned to the Camelina reference genomes (v.2: https://cruciferseq.ca/Csativa_download) using the split-read aligner BWA-MEM (version 0.7.17) with the recommended parameter settings (-T 19) as outlined in the CIRI2 usage instructions. Aligned reads were used as input along with the Camelina reference genome and corresponding genome annotation file for running CIRI2.

CIRI2 and CLEAR pipelines have different filtering criteria for reporting circRNAs. For example, CIRI2 reports intergenic circRNAs, and whereas CLEAR excludes intergenic regions and while CLEAR reports circRNAs with any number of supporting BSJ reads, CIRI2 only reports circRNAs with at least two supporting reads. CircRNA BSJ coordinates may not be entirely accurate as they are dependent on the sequence at the splice junctions and the annotation system of the different pipelines. If small portions of identical sequences exist on both sides of the BSJ, its coordinates can be assigned differently depending on the software and exact supporting reads. To account for this, we combined circRNAs with very similar BSJs (Manhattan distance ≤10 across start and end coordinates) as identical circRNAs. (Supplementary Table S3).

In all treatment samples combined, the CIRI2 pipeline identified a total of 50,922 BSJ reads (>1 read/BSJ) from putative circRNAs in all samples while CLEAR identified 12,843 putative circRNAs based on their BSJ reads (Table 1; Supplementary Table S3). The vast majority of BSJ reads identified by CIRI2 (32,065) originated from 307 distinct intergenic loci. The genic BSJ reads identified by CIRI2 corresponded to 282 exonic and 48 intronic putative circRNAs with loci originating from 256 and 42 unique gene IDs, respectively. There was no overlap in gene IDs between intronic and exonic circRNAs identified by CIRI2.

**TABLE 1:** Comparison of circRNA identified via CIRI2 or CLEAR pipelines. Outputs from CIRI2 (A) and CLEAR (B) were combined to identify all putative circRNA (C). Only 12% of all circRNAs were identified by both pipelines (D). For the CLEAR pipeline, numbers in parenthesis are the number of circRNA BSJ with >1 read.

|  | A CIRI2 |  |  |  | B CLEAR |  |  |
| --- | --- | --- | --- | --- | --- | --- | --- |
|  | total | exonic | intronic | intergenic | total | exonic | intronic |
| Total circRNAs (BSJ reads) | 50,922 | 3,913 | 14,944 | 32,065 | 12,843 (8,937) | 12,629 (8,875) | 214 (62) |
| Total unique circRNAs | 637 | 282 | 48 | 307 | 3239 (775) | 3092 (749) | 147 (26) |
| Total unique cognate gene IDs | 293 | 256 | 42 | NA | 2612 (647) | 2481 (622) | - |
|  | C Combined circRNAs (identified by both and either method) |  |  | D Method comparisons |  |  |  |
|  | total | exonic | intronic | CIRI only | both | CLEAR only |  |
| Total unique circRNAs | 3,447 (1007) | 3,253 (934) | 194 (73) | 208 | 122 (98) | 3,117 (677) |  |
| Total unique cognate gene IDs | 2,763 (843) | 2,602 (781) | 185 (67) | 151 | 146 (101) | 2,466 (546) |  |

The CLEAR pipeline identified a total of 3239 unique circRNAs that originated from 2,612 transcripts with unique gene IDs (Table 1B). Of those, 3092 unique exonic circRNA and 147 unique intronic circRNAs originated from 2481 and 147 unique gene IDs, respectively, with 17 unique gene IDs generating both, exonic and intronic circRNAs. When single-read BSJs were removed from the CLEAR output to identify and compare it to the CIRI2 analysis which has a >1 cutoff, (Table 1; numbers in parenthesis), CLEAR identified 749 unique exonic and 147 unique intronic putative circRNAs originating from 622 and 26 gene IDs respectively, with two genes containing both, an exonic and an intronic putative circRNA.

Combining the outputs of both analysis pipelines, we identified a total of 3,447 unique circRNAs originating from 2,763 genes (Table 1C). Of those 3,447 circRNAs, 208 were only identified by CIRI2 and 3,117 were only identified by CLEAR (Table 1C). Only 122 (12% of total) circRNAs were identified by both pipelines. Those 122 circRNAs that were identified by both pipelines, were only sometimes identified in all samples. For example, circCsa11g034060_1 (Chr11: 15840897-15842933) from Csa11g034060 encoding TOPLESS-related 2, a transcriptional co-regulator in autoimmunity and plant development, was found by the CLEAR pipeline in all 15 samples, while the CIRI2 pipeline only identified this circRNA in 4 of the 15 samples. In some cases, CIRI2 identified circCsa17g055770_1 [Chr17: 19864215-19864901 from Csa17g055770-Suppressor of Auxin Resistance-1 (CsSAR1)] in 8 of the 15 samples, while CLEAR only identified this circRNA in one sample (Supplementary Table S3).

While the computational identification of circRNA by both analysis pipelines provides an apparent additional level of confidence in the detection, we have previously shown that circRNAs identified by only one of those methods, CIRI or CLEAR can be validated using PCR (Budnick et al., 2024). Therefore, we used the results from both pipelines, including those with only 1 read in CLEAR, for a combined total of 3,447 unique circRNAs from 2,763 different cognate genes for further characterization of chromosomal distribution, alternative splice forms, and relative abundances functional characteristics.

Due to the low abundance of circRNA, sequencing depth (reads per sample) could affect the number of circRNAs identified in a sample such that deeper sequencing should yield a higher number of new circRNA up to a saturation point. To investigate whether differences in sequencing depth might account for differences in detection or lack of detection of circRNA, we compared our read depths to the number of total circRNA reads for each sample from both analysis pipelines (Table SI, Figure 2 A, B). While the sequencing depth varied between 43 million and 83 million PE reads per sample, the number of unique circRNAs per million reads in a sample varied from 201 to 521 (CLEAR) and 30 to 61 (CIRI2). The higher number of circRNAs per sample in the CLEAR pipeline is due to the cut-off of BSJ>1 read in CIRI2. We also removed for this analysis a likely mis-annotated light-specific circRNA (*Plant defensin 1.3*, Supplementary Table S3) that was only identified by the CIRI2 pipeline, because it accounted for up to 90% of total BSJ reads in those samples. Blasting the BSJ sequence (100bp) of this putative *defensin 1.3* against the camelina genome identified another genomic area with high sequence identity. We therefore excluded these reads for the subsequent analyses. Sample S17 was removed as an outlier because the number of identified BSJ reads in it was more than an order of magnitude lower than all other samples (Supplementary Table S3). Correlation between sequencing depth (total reads) versus total circRNA reads or unique circRNAs showed no correlation (Figure 2A, B). On the other hand, samples from the same treatment groups clustered together, indicating that sequencing depth was sufficient to saturate the identification of circRNAs and that treatment correlated more with the discovery of circRNAs or circRNA total reads than sequencing depth.

**Figure 2:**
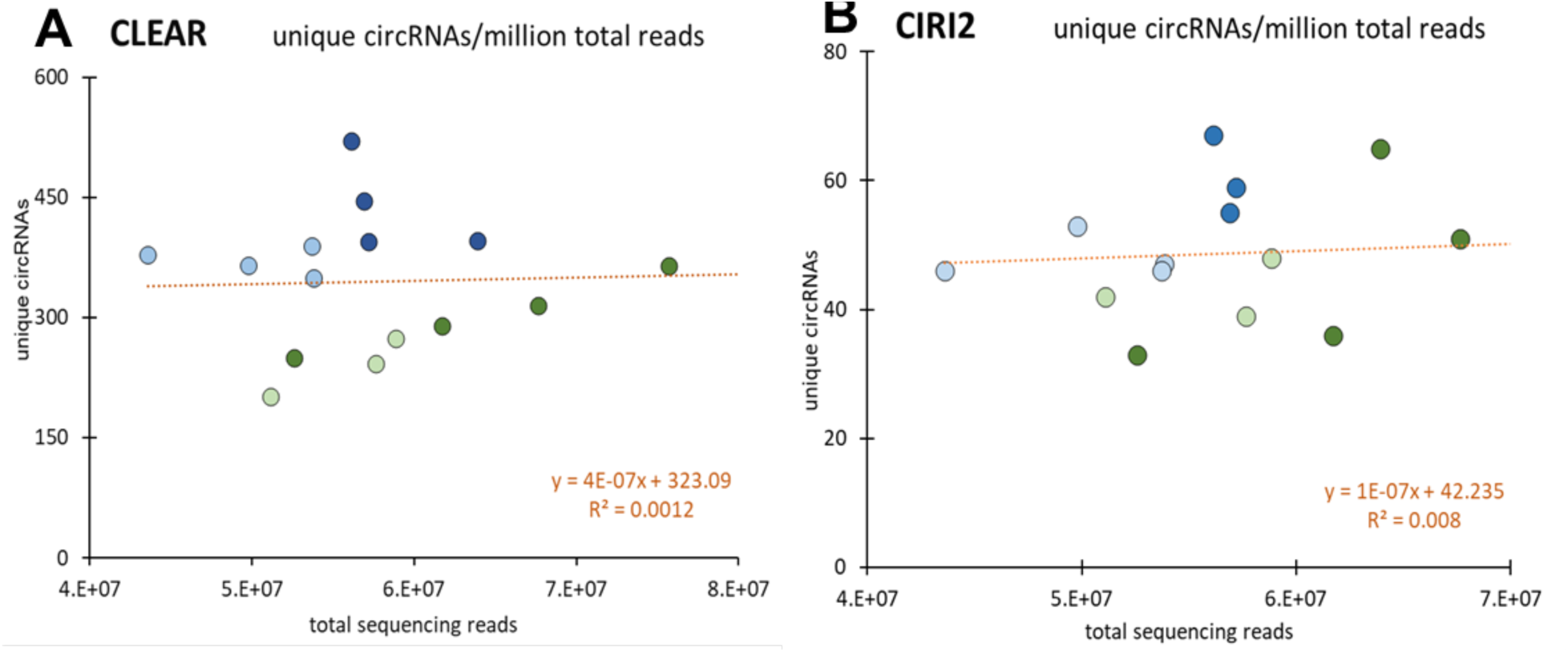
Comparison of read depth and circRNAs. per sample as identified by either CLEAR (A) or CIRI2 (B) shows that there is no significant correlation between sequencing depth to circRNA discovery. Samples from different treatments are colored the same (5L, light green; 5D, light blue; 7L dark green; 7D, dark blue). Supplementary Table S3.

### 3.3. CircRNA can originate from different and multiple homeologs

Genome sequencing of the allohexaploid *Camelina sativa* has shown extensive synteny and colinearity of the camelina subgenomes G1, G2, and G3 with *Arabidopsis thaliana* and *Arabidopsis lyrata* (Kagale et al., 2014). We used the synteny matrix of Arabidopsis and Camelina orthologs (Kagale et al., 2014) to identify and distinguish circRNAs originating from the homeologs from the different subgenomes (Figure 3). Overall, each subgenome had a similar number of genes generating circRNA: 1,072 circRNAs from G1, 1,078 circRNAs from G2, and 1,086 circRNAs from G3. Chromosome 11 in subgenome G1 had about twice as many genes generating circRNA compared to all of the other chromosomes, but Chr11 is the largest chromosome with colinearity to both arabidopsis chromosomes 7 and 8, representing a “fusion” of Cs chr18 and Cs chr10 in the subgenome G2. Overall, the number of circRNA cognate genes were fairly evenly distributed across chromosomes and subgenomes. The lowest percentage of circRNA generating genes was 3.7% from Chr04 and the highest percentage was 5.4% originating from genes in Chr01.

**Figure 3:**
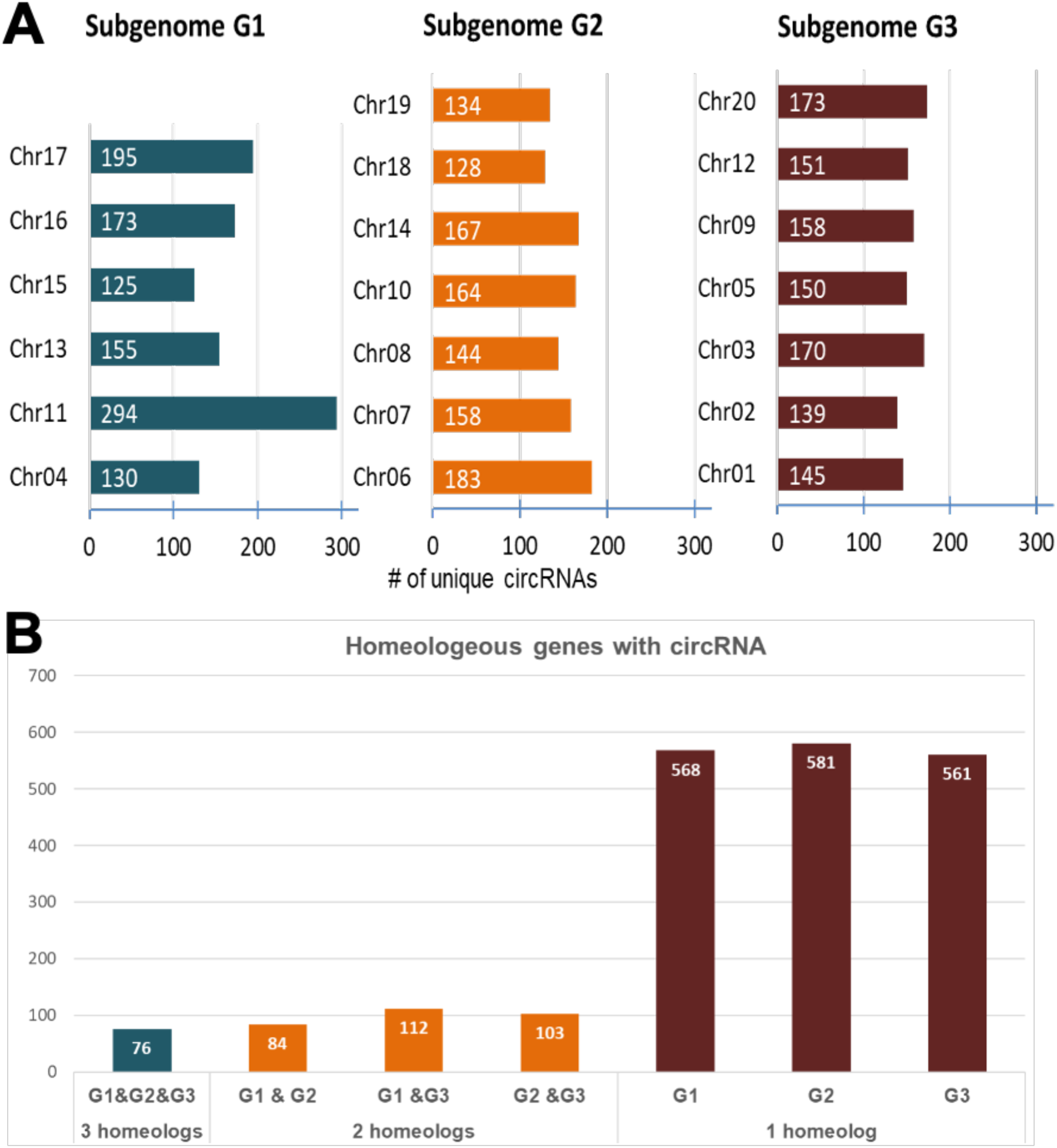
Unique circRNAs from each chromosome (A) organized by subgenome. Each subgenome has about the same number of genes with circRNAs. Most circRNA generating genes originated only from a single subgenome, but 299 circRNAs originated from homeologs from 2 different subgenomes while only 76 circRNAs originated from the homeologs from all 3 subgenomes (B).

While circRNA identified in this study originated to a similar extent from genes from all chromosomes in the subgenomes, most of the circRNAs detected did not originate from all 3 homeologs of their cognate genes (Figure 3B). It is possible, however, that the high sequence identity between some homeologs prevented homeolog distinction of the short BSJ reads.

### 3.4. Constitutive and Treatment-responsive circRNAs

CircRNA identified by both pipelines were analyzed for their general conserved or treatment-specific qualitative and quantitative occurances. Significantly fewer unique circRNAs were identified in the light treatment samples (5L; 7L) compared to the dark treatment samples (5D; 7D) (Figure 4A). There were also significantly more unique circRNAs identified by both pipelines between the 5D and 7D treatment, while there was no significant difference in the number of unique circRNAs between the 5- and 7-day light samples (5L vs 7L). The total number of BSJ reads was higher in the dark vs light treatment samples from either analysis pipeline. There was no significant difference between the dark treatments (Figure 4C, D). This indicates that, while fewer unique circRNAs were expressed in the 5-day samples compared to the 7-day samples, some of those circRNAs were expressed in higher abundance as quantified by total BSJ reads (Supplementary Table S3).

**Figure 4:**
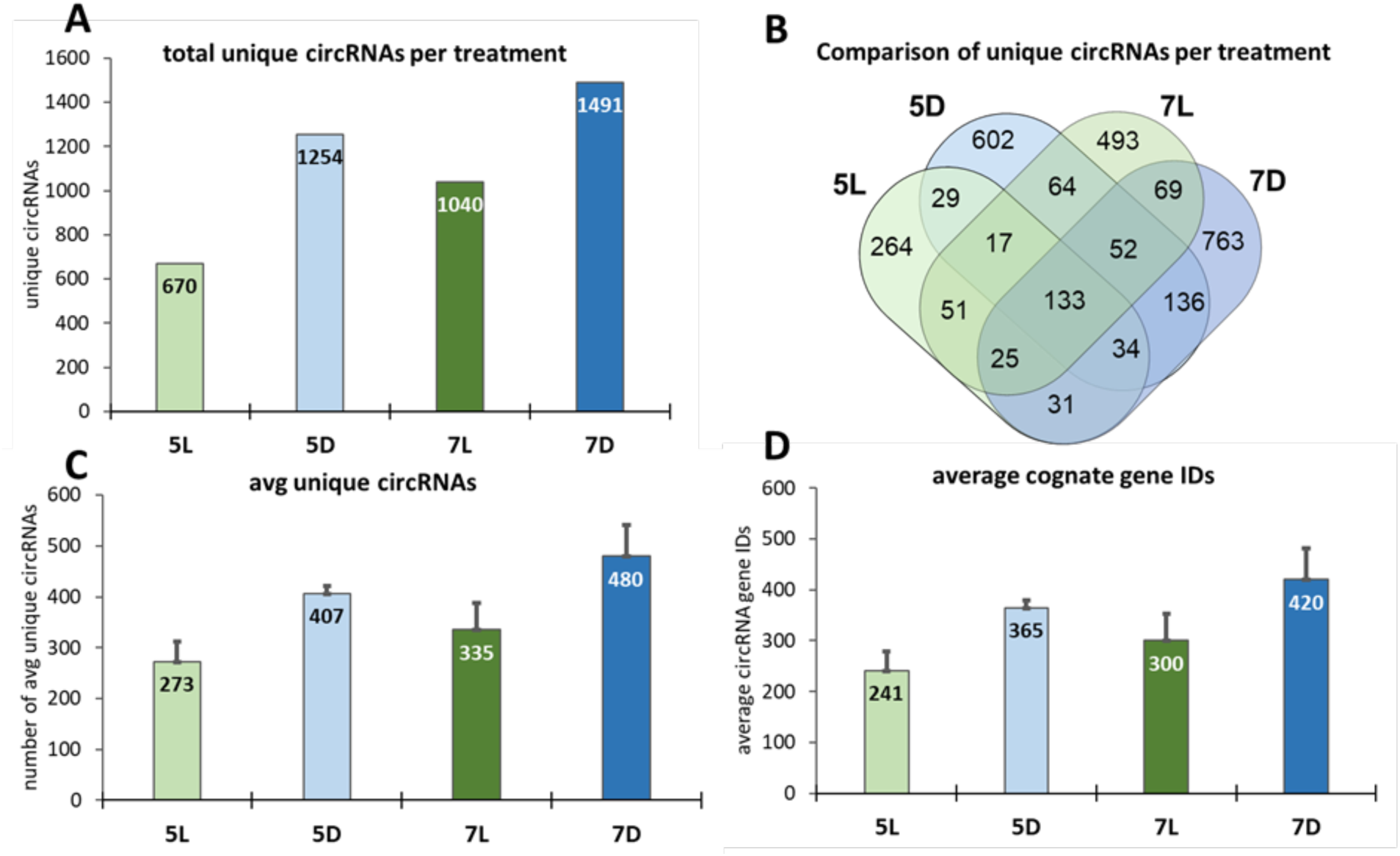
The qualitative and quantitative abundance of circRNAs and their cognate gene IDs varies with treatment. The average number of unique circRNAs identified in the dark-treated seedlings was significantly higher compared to the light-grown seedlings after both 5 days and 7 days of treatment (A). The same pattern was identified for the average unique cognate genes that those circRNA’s derived from (B). The average total number BSJ reads was significantly higher in the dark-treated samples as well, but there was no significant difference between the samples after 5 vs 7 days of treatment (C, D). For data see Supplementary Table S3.

We next analyzed circRNAs to identify constitutively expressed ones within a treatment or across all treatments to serve as “house-keeping” circRNAs in quantitative comparisons. Given the low abundances of some circRNAs and differences in the analysis pipelines, we termed unique circRNAs that were present in ≥2 samples of a treatment as “conserved” in that treatment, and those that were present in all samples of a treatment were termed “strictly conserved”; Figure 5; Supplementary Table S3).

**Figure 5:**
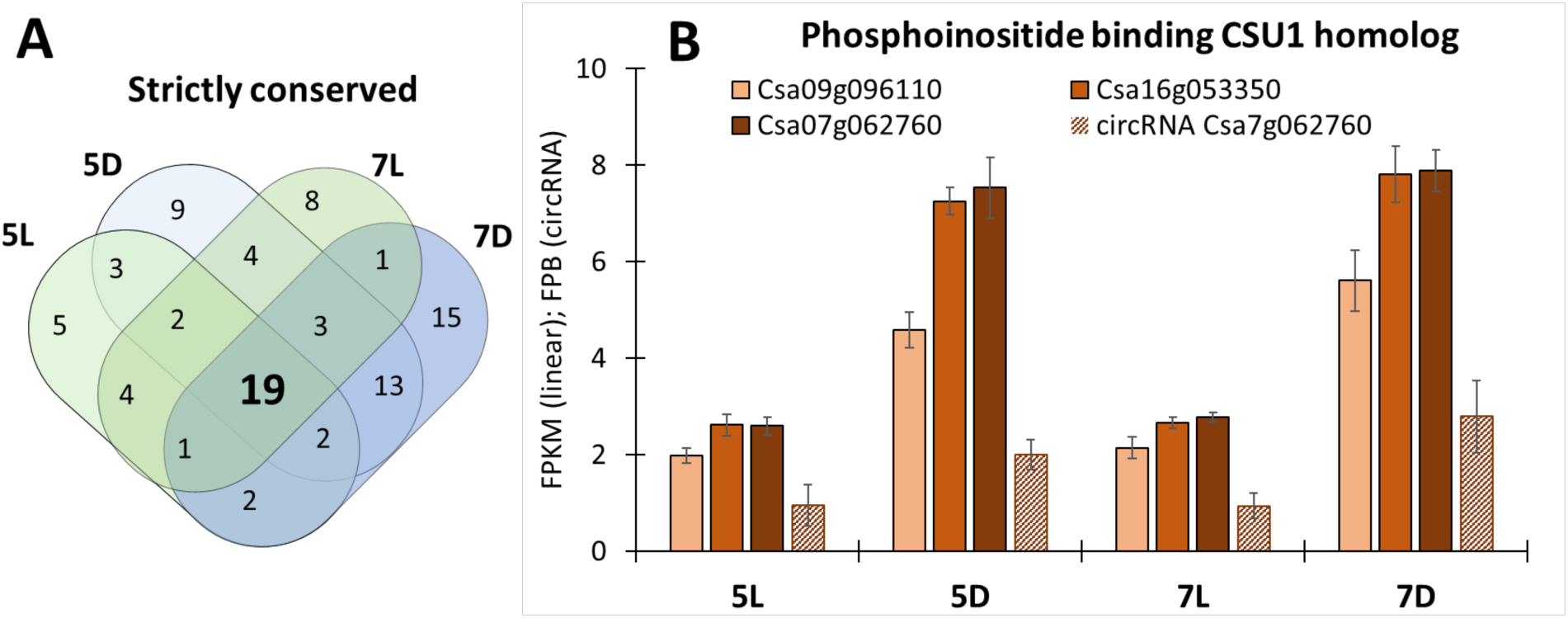
Comparison of circRNAs expressed under different treatments. We compared clustered, combined circRNAs identified from all samples in a treatment (5L, 5D, 7L, 7D) to identify circRNAs that were strictly conserved among all treatments (A). A group of 19 unique circRNAs (bold) originating from 14 unique gene IDs were present in all samples (Table 2). One example is the *Phoshoinositide binding CSU homolog* circCsa07g062760_1 which is strictly conserved in all samples (B).

**TABLE 2:** “Strictly conserved” circular RNA present in all samples. (19 from Figure 5). These circRNA originated from 16 genes, some of which had more than one circRNA identified in multiple samples (those conserved in all samples are in bold).

| circRNA genome coordinates | Cognate Gene ID | Functional annotation | Arabidopsis ortholog |
| --- | --- | --- | --- |
| **Chr1: 19060177_19061728**<br>Chr1: 19060367_19061984 | Csa01g038070 | *aconitase 3* | AT2G05710.1 |
| **Chr1: 10840971_10842502** | Csa01g025690 | *RNA recognition motif XS domain protein* | AT3G22430 |
| **Chr10: 2977184_2981849**<br>**Chr10: 2978450_2982754** | Csa10g008100 | *poly(A) binding protein 2* | AT4G34110 |
| **Chr11: 15840897_15842933** | Csa11g034060 | *TOPLESS-related 2* | AT3G16830 |
| **Chr12: 31990610_31993495**<br>Chr12: 31990891_31993710 | Csa12g084210 | *Ribulose biphosphate carboxylase (small chain) family protein* | AT5G38430 |
| **Chr15: 21750182_21751375** | Csa15g064470 | *minichromosome maintenance 9* | AT2G14050 |
| **Chr18: 19928903_19929731** | Csa18g039550 | *nitrogen fixation S (NIFS)-like 1* | AT5G65720 |
| **Chr18: 9670136_9676228**<br>Chr18: 9670504_9676638<br>**Chr18: 9672080_9679354** | Csa18g019070 | *ubiquitin-protein ligase 2* | AT1G70320 |
| **Chr2: 2216430_2223563** | Csa02g007690 | *citrate synthase 5* | AT3G60100 |
| **Chr20: 18063735_18065784**<br>**Chr20: 18064891_18065784** | Csa20g052830 | *DEAD box RNA helicase (RH3)* | AT5G26742 |
| **Chr3: 3385824_3387133**<br>Chr3: 3386059_3387427 | Csa03g011570 | *DNA ligase 1* | AT1G08130 |
| **Chr4: 8139584_8141441** | Csa04g016640 | *CLEC16A-like (TT9)* | AT3G28430 |
| **Chr6: 5083634_5085478** | Csa06g009030 | *NADPH:quinone oxidoreductase* | AT3G27890 |
| **Chr7: 31836782_31837722** | Csa07g062760 | *phosphoinositide binding Ring finger E3-ligase CoSupp. of COP1 (CSU1)* | AT1G61620 |
| **Chr9:27331656_27332922** | Csa09g071160 | *DNase I-like family protein* | AT3G58560 |
| **Chr1:15332231_15333347** | Csa01g032940 | *lncRNA171, put. trichohyalin-like* | Camelina specific |

Comparison of these treatments “conserved” circRNA showed that 19 circRNAs were present in all samples tested (Figure 5A). We consider these 19 as “housekeeping”circRNAs based on their presence in all samples (Table 2).

Those 19 “housekeeping” circRNA gene IDs present in all samples included the RUBISCO SSU (Csa12g084210), Ubiquitin-protein ligase 2 (Csa18g019070), DEAD box RNA helicase (RH3) (Csa20g052830), TOPLESS-related 2 (Csa11g034060), DNA ligase 1 (Csa03g011570), aconitase 3 (Csa01g038070), CLEC16A-like protein (Transparent Testa 9 homolog; Csa04g016640), nitrogen fixation S (NIFS)-like 1 (Csa18g039550), citrate synthase 5 (Csa02g007690) and NADPH:quinone oxidoreductase (Csa06g009030), as well as phosphoinositide binding CoSuppressor of COP1 (CSU1; Csa07g062760, Figure 5B). CSU1 is an example of a gene where all 3 homeologs were expressed and their linear transcript abundances were all responsive to L/D treatment, but only one homeolog, Csa07g062760, was identified as the cognate gene for a circRNA (Table 2, Figure 5B).

In contrast to circRNAs that were identified as conserved in all samples, we identified circRNAs that were specific to a treatment. Treatment-specific circRNAs were identified through a comparison of circRNAs that were present in a majority of samples for one treatment (“treatment-conserved”) but not present in any samples of the other treatments (Figure 6A-D). CircRNAs that were identified as specific to a treatment are listed in Table 3.

**Figure 6:**
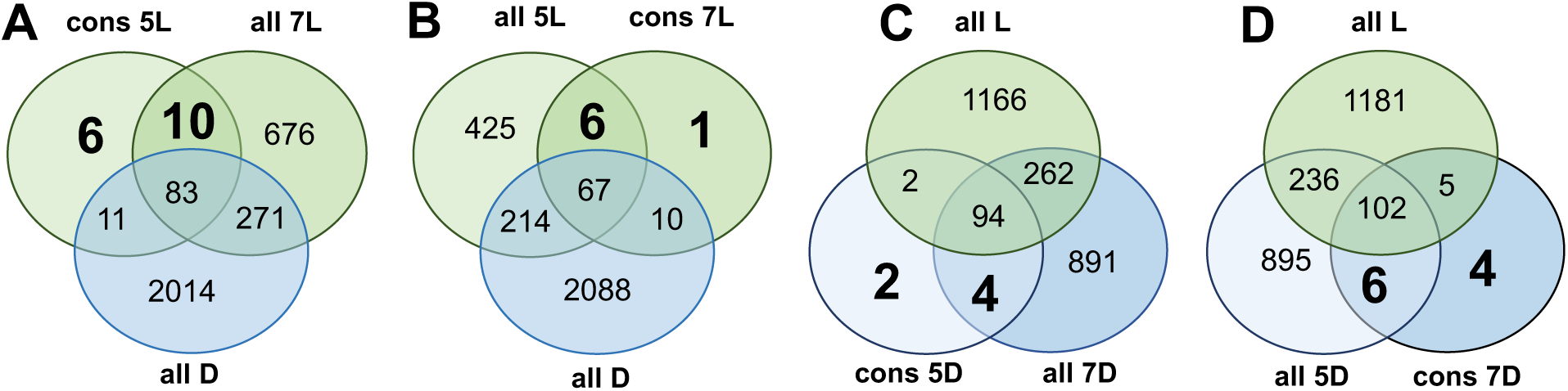
Conserved, treatment-specific circRNAs. Unique circRNAs that were present in most samples of a treatment (cons) were compared to those in all other treatments to identify treatment-specific circRNAs from 5L (A), 7L (B), 5D (C), and 7D (D). Treatment-specific circRNAs (bold numbers) are listed in Table 3.

**Table 3:**
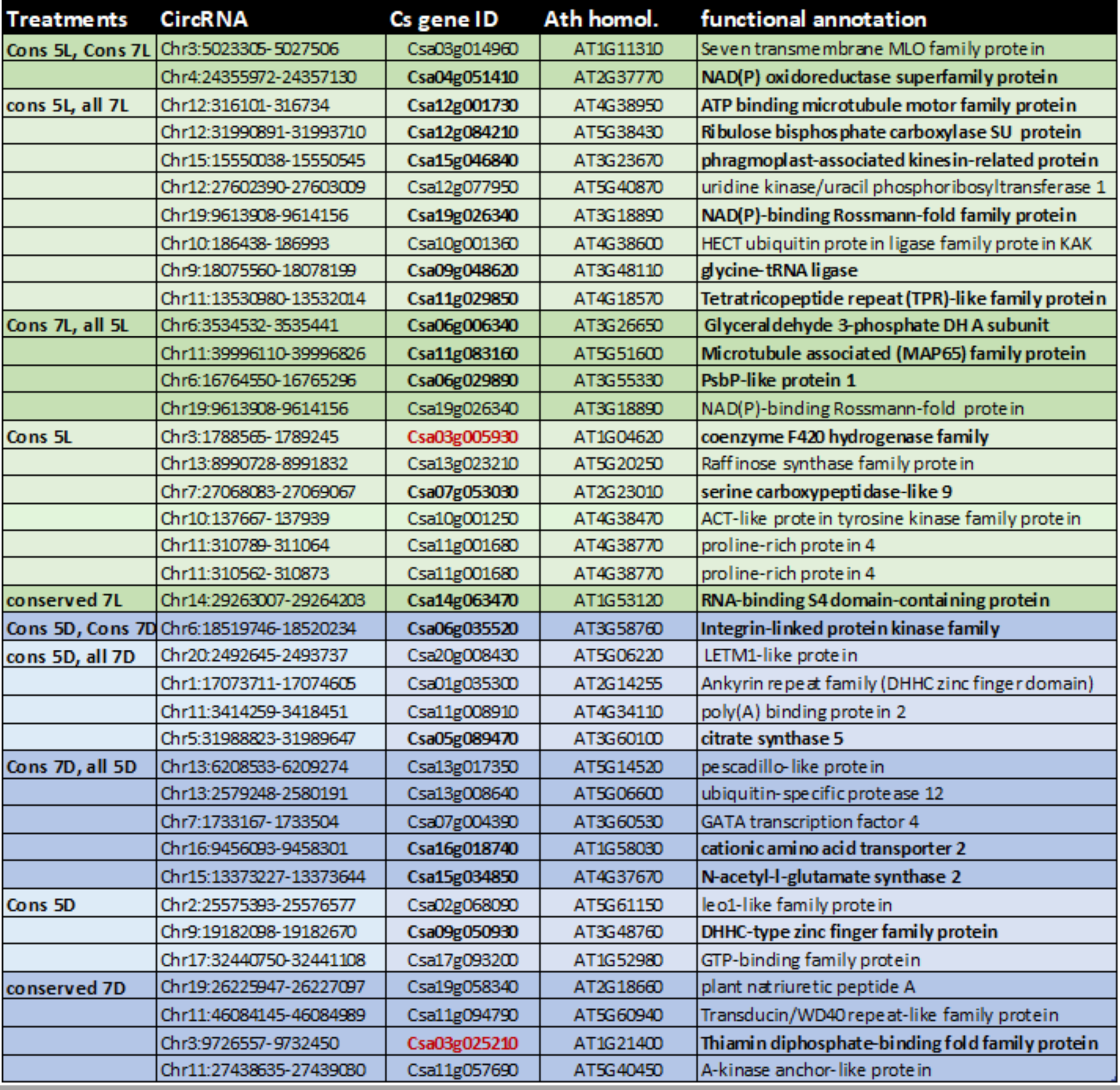
Treatment-specific circRNA. Conserved (cons) circRNAs from a treatment were compared to all circRNAs of the other treatments (Figure 6 A-D). The treatment-specific circRNAs (Figure 6, bold) are listed in this table. Bolded circRNA gene IDs refer to those circRNAs that originate from linear DEGs (Figure 7).

Two circRNAs were conserved in both 5L and 7L treatments and absent from dark treatments. These two circRNAs originated from Csa03g014960, a “Seven transmembrane MLO family” protein gene and Csa04g051410, an “NAD(P) oxidoreductase superfamily protein” gene (Figure 8A, Table 3). The Seven transmembrane MLO family is a large family of membrane proteins of which members have been shown to confer broad resistance to fungal pathogens (Büschges et al., 1997). The NAD(P) oxidoreductase is orthologous to the Arabidopsis AKR4C9, a nuclear encoded, plastid-localized aldo-keto reductase with broad substrate specificity that is responsive to abiotic stresses including cold, salt and dehydration (Simpson et al., 2009).

Only one circRNA was conserved in both dark treatments but absent in all light treatments. This single circRNA is circCsa06g035520_1 originating from Csa06g035520, a gene annotated as Integrin-linked protein kinase family gene that has been shown to be a link between plant defense pathways and K+ homeostasis (Figure 8B). Integrin-linked protein kinases are involved in signal transduction and can regulate the plant’s response and sensitivity to osmotic stress, nutrient availability, and biotic interactions (Brauer et al., 2016). These kinases are composed of Ankyrin repeats and phosphorylate MAPkinases as part of their signal amplification/transduction (Popescu et al., 2017).

CircRNA from several key enzymes of the Calvin-Benson Cycle, including Ribulose 1,5 bisphosphate carboxylase/oxygenase small subunit (RUBISCO-SU), Glyceraldehyde 3-phosphate dehydrogenase, as well as sugar metabolism genes like Raffinose synthase are specifically expressed under light conditions. Dark-specific circRNAs originated from citrate synthase 5, a key enzyme involved in the Krebs Cycle, which is essential in the dark to generate ATP for metabolic conversions during respiration. Interestingly, a different ortholog of citrate synthase 5 circRNA was identified as a “housekeeping” circRNA (Table 2). Dark-specific synthesis of circRNA was also detected for amino acid metabolism with the cationic amino acid transporter 2 (Csa16018740) and the N-acetyl-glutamate synthase 2 (Csa15g034850), which is essential for arginine biosynthesis. While the circRNAs identified in Table 3 were detected only under a specific treatment, linear RNA cognates for most of these genes were present in all or most treatments and many of those showed treatment-specific changes in transcript abundances (Figure 8).

### 3.5. CircRNAs from Differentially Expressed Genes (DEGs)

CircRNA has been shown to regulate transcript abundances via different mechanisms including increased transcription rates or transcript stability via miRNA sequestration. We analyzed the correlation of treatment-specific changes in linear RNA abundances with total circRNA occurrence by comparing linear transcripts from differentially expressed genes (DEGs) with all circRNAs from those treatments (Figure 7A) or with only the treatment-specific circRNAs identified in Table 3 (Figure 7B).

**Figure 7:**
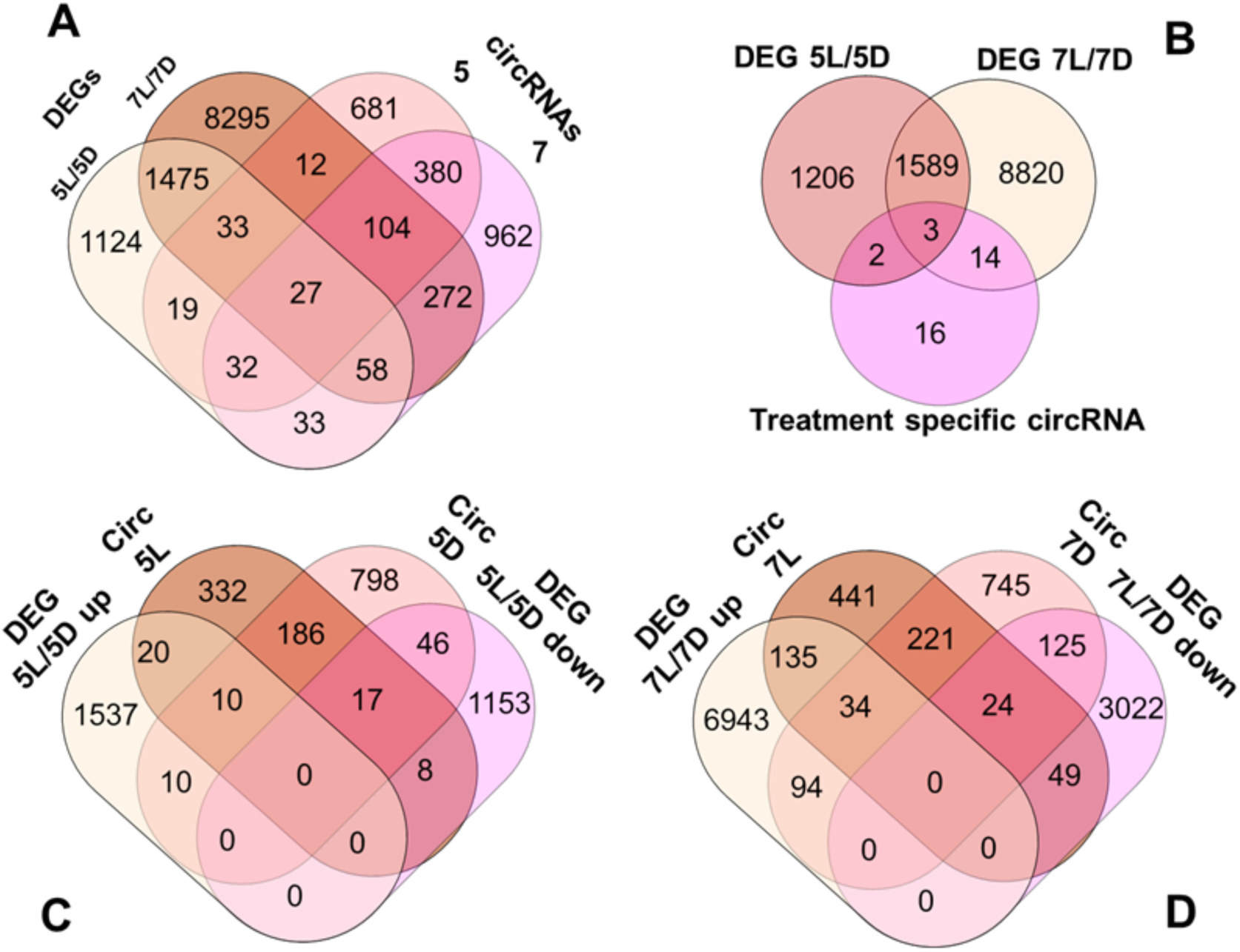
Treatment-specific DEG linear transcripts and their circRNAs. DEGs contrasting L vs D treatments after 5 and 7 days against all circRNAs (A). Only 19 of the conserved and treatment-specific circRNAs as identified in Table 3, originated from DEGs (B). Significantly up or down regulated DEGs per treatment were compared to circRNA at either light or dark treatment after 5 days (C) or 7 days (D).

Linear transcript abundances were normalized for differences in library size, and transcript abundance was expressed as reads counts per million mapped reads (cpm). Reads corresponding to intergenic regions and transcripts with ≤ 10 reads were discarded for this analysis. A total of 73,234 unique transcribed genes were identified and their abundance normalized (Supplementary Table S4). With 2,763 circRNA generating transcripts, the overall percentage of transcribed genes generating circRNA is 3.8%.

Transcripts with significantly different abundances (DEGs, FDR≤ 5*10^−4^) between light and dark treatments after 5 or 7 days of growth (Supplementary Table S4) were compared to all circRNA gene IDs identified in the 5 day and 7 day treatments. We identified 2,801 DEGs after 5 days of treatment (5L vs 5D) and 10,426 DEGs after 7 days of treatment (7L vs 7D) with significant differences in transcript abundances (Figure 7).

Overall, 2,801 unique transcripts were differentially expressed between 5L and 5D treatments (DEGs 5L/5D), while 10,426 transcripts were differentially expressed between light and dark treatments after 7 days of growth (DEGs 7L/7D). Between both contrasts, 1,593 genes were differentially expressed at 5 and 7 L/D contrasts. These DEGs were compared to 1,438 unique circRNA cognate gene IDs identified after 5 day treatments and 1,868 unique circRNA cognate gene IDs from the 7 day treatment (Figure 7A). Overlap between the categories showed that 111 of the 2,801 DEGs from 5L/5D had corresponding circRNAs and 461 of the 10,426 DEGs in the 7 day treatment were correlated with the formation of circRNA, accounting for 3.96% and 4.42% of DEGs with circRNA respectively. A similar comparison of circRNA from either both 5 day or 7 day treatments with significantly up or down-regulated DEGs of the respective treatments (Figures 7B and 7C) showed that 2.5% of 5L/D up-regulated DEGs (higher in light grown than in dark grown seedlings) were cognate genes forming circRNAs, while 5.8% of DEGs that were higher expressed after 5 days in the dark generated circRNA - about twice as many. A similar result came from the same comparison after 7 day treatments, with 3.7% of DEGs with higher abundance in the light producing circRNA and 6.1% of those DEGs with higher abundance in the dark-grown seedlings.

In a more stringent comparison, we only compared circRNA from genes that were already identified as conserved within a treatment (from Table 3) to those L or D specific DEGs (Figure 7B). This comparison identified 2 conserved and treatment-specific circRNAs from the 5L/5D comparison and 14 circRNAs from the 7L/7D DEGs, and 3 treatment-specific circRNAs with their cognate linear transcripts showing significant differential expression at both L vs D comparisons. The linear DEGs with treatment-specific circRNAs are reported in Table 3.

Among the genes with treatment-specific circRNA and differentially expressed linear transcripts we identified one specific homeolog to each of the RUBISCO SSU (Csa12g084210). Two treatment-specific, conserved circRNAs showed differential expression of their linear cognates and had higher expression in the dark: the transcript for a coenzyme F420 hydrogenase genes (Csa03g005930) as well as the a Thiamin diphosphate-binding fold family protein (Csa03g025210) (shown in bold and red, Table 3). The coenzyme F420 hydrogenase gene transcript is particularly interesting because its circRNA contains an intronic region from the cognate gene and is correlated with an increase in abundance of an intron-retaining alternatively spliced version of the cognate linear transcript and a corresponding decrease in the full-length exact match intron chain transcript (see Figure 9B). The Arabidopsis ortholog is identified as the 7-hydroxymethyl chlorophyll a (HMChl) reductase that catalyzes the last step in chlorophyll a biosynthesis (Meguro et al., 2011). A homolog of the same gene in Arabidopsis, called Thiamin diphosphate-binding fold family protein, is a branched chain alpha-keto acid dehydrogenase (E1 alpha subunit) which is involved in catabolic processes especially under sucrose starvation and in the dark (Peng et al., 2015). These findings suggest that this enzyme involved in protein breakdown in the etiolated seedlings of Camelina.

As examples for treatment-specific circRNA and their linear cognate DEGs, the ratio of circRNA counts corresponding to the BSJ to linear RNA (FPBcirc; calculacted in the CLEAR pipeline (Ma et al., 2019) are shown below for the light-specific circCsa03g014960_1 (seven transmembrane MLO family protein), the dark-specific circRNAs circCsa06g035520_1 (Integrin-linked protein kinase family) and circCsa05g089470_3 (Citrate Synthase 5). In addition to the dark-specific circRNA, a different homeolog of Citrate Synthase 5 (Csa02g007690) also generates a circRNA and although it is found in all treatments, it increases in abundance in the dark treated samples. The opposite response was found in the Rubisco small subunit genes. One of the three homeologs of the Rubisco small subunit (Csa12g084200) generated a circRNA that was present in all samples regardless of treatment but was found in higher abundance in the light treated samples (Figures 8 A-C).

**Figure 8:**
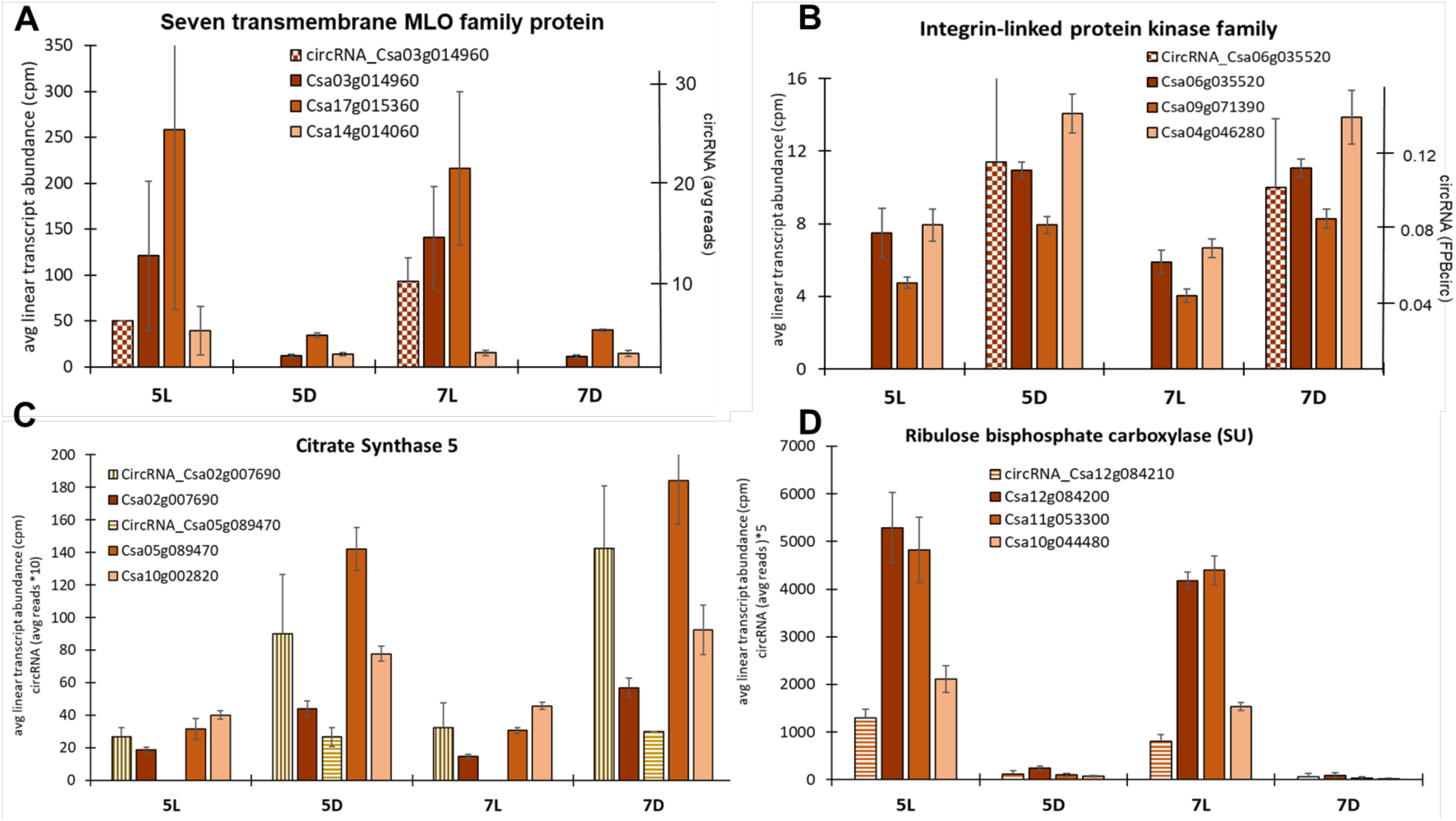
Treatment-specific circRNA abundances compared to their linear DEG cognate transcript abundances. A “Seven transmembrane MLO family protein” transcript generates a light-specific circCsa03g014960_1 with circRNA only originating from one homeolog, but all 3 homeologs have LD treatment responses (A). The higher abundance in the etiolated samples 5D and 7D occurs for the dark-specific circRNA of circCsa06g035520_1 Integrin-linked protein kinase family transcript (B). One homeolog of Citrate Synthase 5 (Csa5g089470) (C) as well as one RUBISCO SSU homeolog (Csa12g084210) (D) are present in all treatments but show higher abundances in response to dark or light, respectively. Another Citrate synthase homeolog (Csa5g089470) generated circCsa05g089470_3 that was only detectable under dark treatment (C). Abundances of circRNAs were so low that they were displayed on a secondary axis (A, B) or presented as 10x on the primary axis (C, D).

Citrate synthase is a key enzyme in the Krebs cycle and is involved in generating redox equivalents and ATP from catabolic reactions. It is also important in providing precursors for lipid and amino acid biosynthesis in plants (Y. Zhang & Fernie, 2023). Rubisco on the other hand is the key enzyme in photosynthetic CO_2_ fixation. This explains the higher abundance of Rubisco transcripts in the light grown seedlings compared to the higher abundance of Citrate synthase transcripts in the etiolated, dark-grown seedlings.

### 3.6. Alternative splice variants of linear RNA correlate with circRNA presence

One of the functions shown for circRNA in plants is the formation of R-loops (RNA:DNA hybrids) that result in the formation of alternatively spliced transcripts and their protein products (Conn et al., 2017). We, therefore, performed a splice-aware alignment and assembly of the sequencing reads to identify different splice versions of linear genic RNAs from our transcriptome data and compared their abundances to the presence of circRNAs (Table II).

Fragments that aligned to the Camelina reference genome and resulted in distinct, primary alignments were used as input for transcript assembly using StringTie (v 2.2.1) (M. Pertea et al., 2016). Initial transcript assembly was performed for each sample separately based on HISAT2 read alignments using known Camelina genome annotations (https://cruciferseq.ca/Csativa_download) as a guide. Assemblies for all samples were merged into a single assembly containing all known transcripts, novel transcripts, and transcripts with retained introns. Transcript abundances were then re-estimated for each sample using the final, merged assembly as the reference genome as a guide. Attributes of novel transcripts were compared to the known, annotated transcripts using GFFcompare in the GFF Utilities package (G. Pertea & Pertea, 2020).

Overall, the final splice-aware assembly resulted in 139,896 transcripts of which 133,803 originated from genes and 6,093 were mapped to intergenic regions. A total of 75,753 genic linear transcripts matched the annotated gene exon-intron structure, identified with (=) in Table 4. (Supplementary Table S5). A total of 22,587 of those complete genic transcripts also expressed alternative splice versions of the transcript, while 53,166 full length genic transcripts did not have any alternatively spliced transcripts detected (Figure 9). In a first comparison with all identified intronic and exonic circRNAs, 1.8% (928) of transcripts that were only identified as complete without AS versions had circRNA, while 7.8% of complete (=) transcripts that had also at least one AS version transcribed and also served as cognate transcripts for circRNA. Among those transcripts that were only identified as AS versions, with no full length (=) transcript observed in our dataset, 4.6% served as cognate transcripts of identified circRNAs. This correlation indicates a potential connection between circRNA biogenesis and alternative splicing events - a correlation that seems obvious due to the need of the splicing machinery for circRNA production, but also due to the functional identification of R-loops involved in exon skipping (Conn et al., 2017; X. Liu et al., 2021).

**Figure 9:**
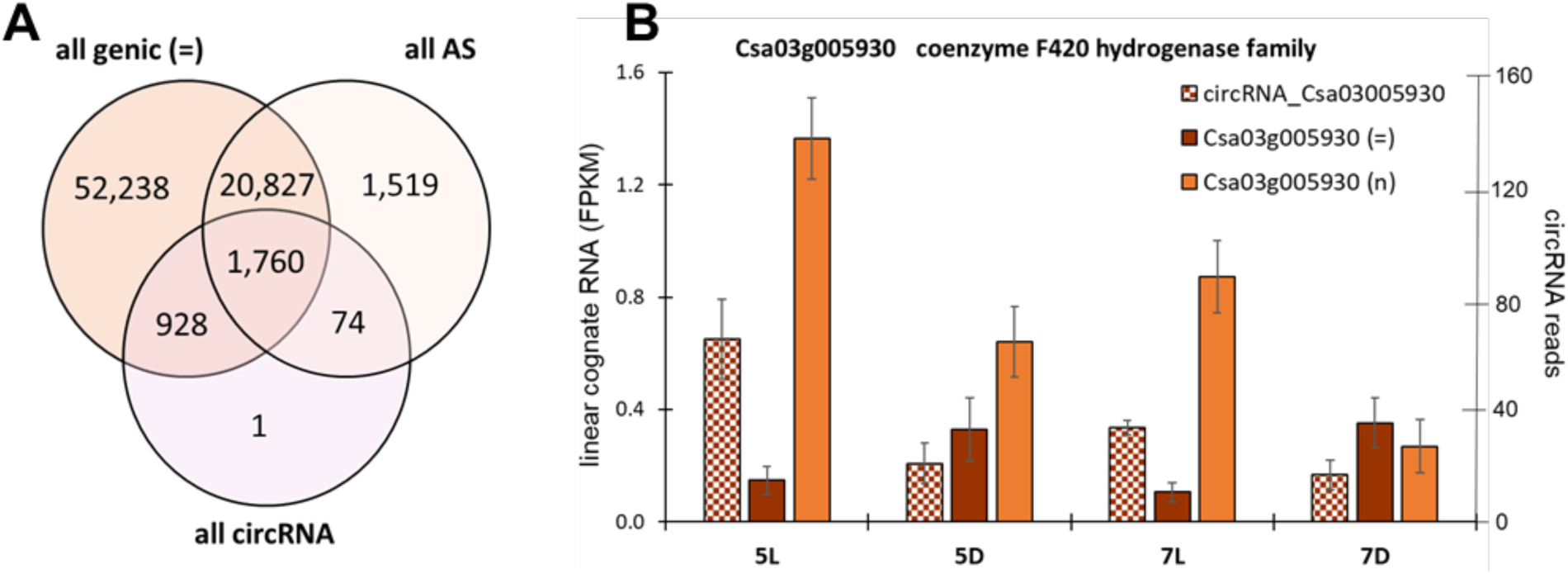
Venn diagram comparing full genic transcripts (all genic (=)) with all alternative splice versions (all AS) identified by the splice-aware software and all genic circRNA (A). The treatment-specific response of an intronic circRNA correlates with an intron-retaining splice-variant of the gene Csa03g005930 but is opposite to the full-length exact match transcript (B).

**TABLE 4:**
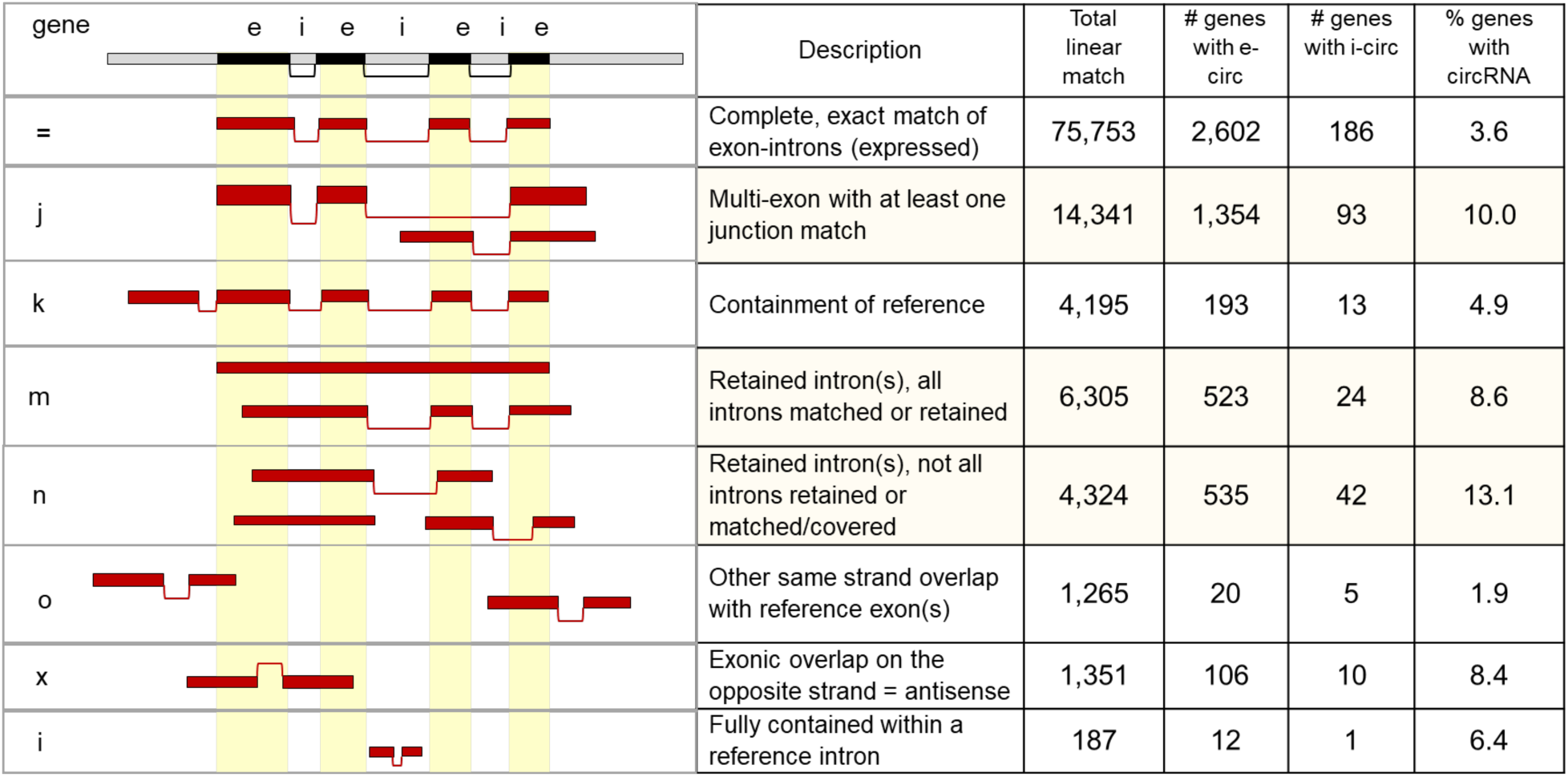
Correlations between different splice variants of genes and their circRNAs. Using splice-aware alignments, we identified for each gene the presence or absence of circRNA in different classes of splice variations (j, k, m, n, o, x, i) based on the genome identification. The default (=) was the exact match of the identified linear genic RNA with all introns removed and all exons present.

| gene | e i e i e i e | Description | Total linear match | # genes with e-circ | # genes with i-circ | % genes with circRNA |
| --- | --- | --- | --- | --- | --- | --- |
| = |  | Complete, exact match of exon-introns (expressed) | 75,753 | 2,602 | 186 | 3.6 |
| j |  | Multi-exon with at least one junction match | 14,341 | 1,354 | 93 | 10.0 |
| k |  | Containment of reference | 4,195 | 193 | 13 | 4.9 |
| m |  | Retained intron(s), all introns matched or retained | 6,305 | 523 | 24 | 8.6 |
| n |  | Retained intron(s), not all introns retained or matched/covered | 4,324 | 535 | 42 | 13.1 |
| o |  | Other same strand overlap with reference exon(s) | 1,265 | 20 | 5 | 1.9 |
| x |  | Exonic overlap on the opposite strand = antisense | 1,351 | 106 | 10 | 8.4 |
| i |  | Fully contained within a reference intron | 187 | 12 | 1 | 6.4 |

Alternative splicing results in different transcript versions with respect to exon and intron content or antisense presence. We analyzed the occurrence of circRNA in different alternative splice categories (Table 4). Among the novel transcripts identified, 14,341 contained a partial match to at least one exon-intron boundary of an annotated transcript (‘j’ classification in Table 4) and retained introns either partway (4,324; category ‘n’) or spanning the full length of the transcript (6,305; category ‘m’). Other attributes identified included 4,195 transcripts containing additional exons (category ‘k’) or 1,265 with additional exons overlapping the same strand (category ‘o’). The remaining transcripts showed exonic overlap with the opposite strand (1,351; category ‘x’) or fully contained features within an intron (187; category ‘i’). Exon skipping or alternative start or end sites were found for 19% of transcripts (category ‘j’), while intron-retention (categories ‘m’ and ‘n’) were identified in 12% of transcripts.

We compared all exonic and intronic circRNA gene IDs to those different AS transcript categories to evaluate if circRNAs are overrepresented in any specific AS category (Table 4). The highest percentage of circRNA was found in genes with exon-skipping events (10%; j) and in intron retention (12%; n and m).

### 3.7. Relative circRNA abundances

Detection of circRNAs is dependent on many aspects including sequencing methods and depth (>60 million reads per sample), quality of RNA extracted, and the quality of the genome sequence and circRNA detection pipelines (see Figure 2). The CLEAR circRNA detection pipeline analyzes not only the number of reads containing a BSJ from a specific circRNA, but also normalizes read abundance as the BSJ Fragments per Billion (FPBcirc) based on the circRNA BSJ reads and a normalizes the abundance of the linear fragments spanning that splice junction (FPB linear) in the same manner. The ratio of FPBcirc and FPBlinear provides a ratio of circRNA to linear RNA, the CircScore. While we will discuss the validity and problems with this analysis, these ratios can indicate a potential function of the circRNA.

The highest read counts (127-283 per sample) originated from Rubisco SSU circRNA. However, due to the high abundance of Rubisco-SSU linear RNA, the circScore (circRNA: linear RNA) was low (ca. 0.01 circRNA:linear RNA). As shown above (Figure 8D), these circRNAs and their cognate linear RNAs were light-responsive.

A total of 53 circRNAs originating from 51 unique gene IDs had read numbers equal to or higher than 10 (283 to 10 reads). All circRNAs identified as strictly conserved circRNAs were found in all samples (Table 2) and had read numbers ≥10, suggesting a bias of “conservation between all samples” and circRNA abundance. High circRNA abundance was also found for the light-specific circRNAs for Csa03g014960 (Seven transmembrane MLO family protein, Figure 8A, circCsa03g014960_1) and Csa03g005930 (coenzyme F420 hydrogenase family, Figure 9B, circCsa03g005930_2). Some of the other circRNAs with high abundance include the nitrate transporter 1.1 (Csa14g015000), a Suppressor of PhyA (SPA1)-related protein (Csa01g017890) from a gene family which has been shown to function redundantly in suppressing photomorphogenesis in dark- and light-grown seedlings. High circRNA read counts were also found for Phospholipase Dbeta 1 (PLDbeta, Csa06g048050) and the Plastid Lipase 3 (PLIP3, Csa05g094920), both of which are involved in biotic responses. Lower read counts (between 2-9 reads) were found for 1,002 unique circRNAs originating from 839 unique gene IDs. As discussed above, CLEAR did not have a cut-off for circRNA BSJ reads, and so identified an additional 2,786 unique circRNAs with only a single read, corresponding to 2,299 unique cognate gene IDs. While some of these may in fact be true circRNAs, they are unlikely to be targets for further investigation without additional read support.

Another important measure is the ratio of circRNA abundance to its linear cognate RNA abundance, the circScore (Supplementary Table S3). The highest circScores (17 to 8) were calculated for 19 circRNAs that only had 1 or 2 BSJ read counts and no reads assigned to the linear transcript in the circRNA analysis pipelines. Only 16 circRNAs had more than 1 BSJ read count and a circScore higher than 1 (more circRNA than linear cognate RNA). The circRNAs with the highest circScores of those 16 were two of the strictly conserved circRNAs from the DNA Ligase 1(Csa03g011570) and CLEC16A-like (TT9; Csa04016640). The other 3 highest circScores with BSJ reads >4 were identified for the Nucleoporin interacting component (Nup93/Nic96-like) family protein (Csa04g043770) involved in mRNA/protein transport across the nuclear membrane (Tamura et al., 2010), a 3-Dehydroquinate synthase (Csa06g010030), catalyzing a reaction of the shikimate pathway for the biosynthesis of aromatic amino acids, and Carbon catabolite repressor 4 (CCR4, Csa09g071160), which encodes a deadenylase involved in mRNA processing and the regulation of carbohydrate metabolism.

One way to categorize circRNA for potential functional involvement is through Gene Ontology (GO) analysis. We use the Panther GeneOntology database (v18) to identify biological processes in which genes containing circRNA are overrepresented in our circRNA pools based on experimental treatments (Mi et al., 2013; Thomas et al., 2022). GO term analysis of Camelina genes was conducted using the biological process GO annotations for orthologous genes in Arabidopsis. We categorized all circRNAs by GO terms for biological processes in order to compare over-represented categories between the treatments (Figure 10).

**Figure 10:**
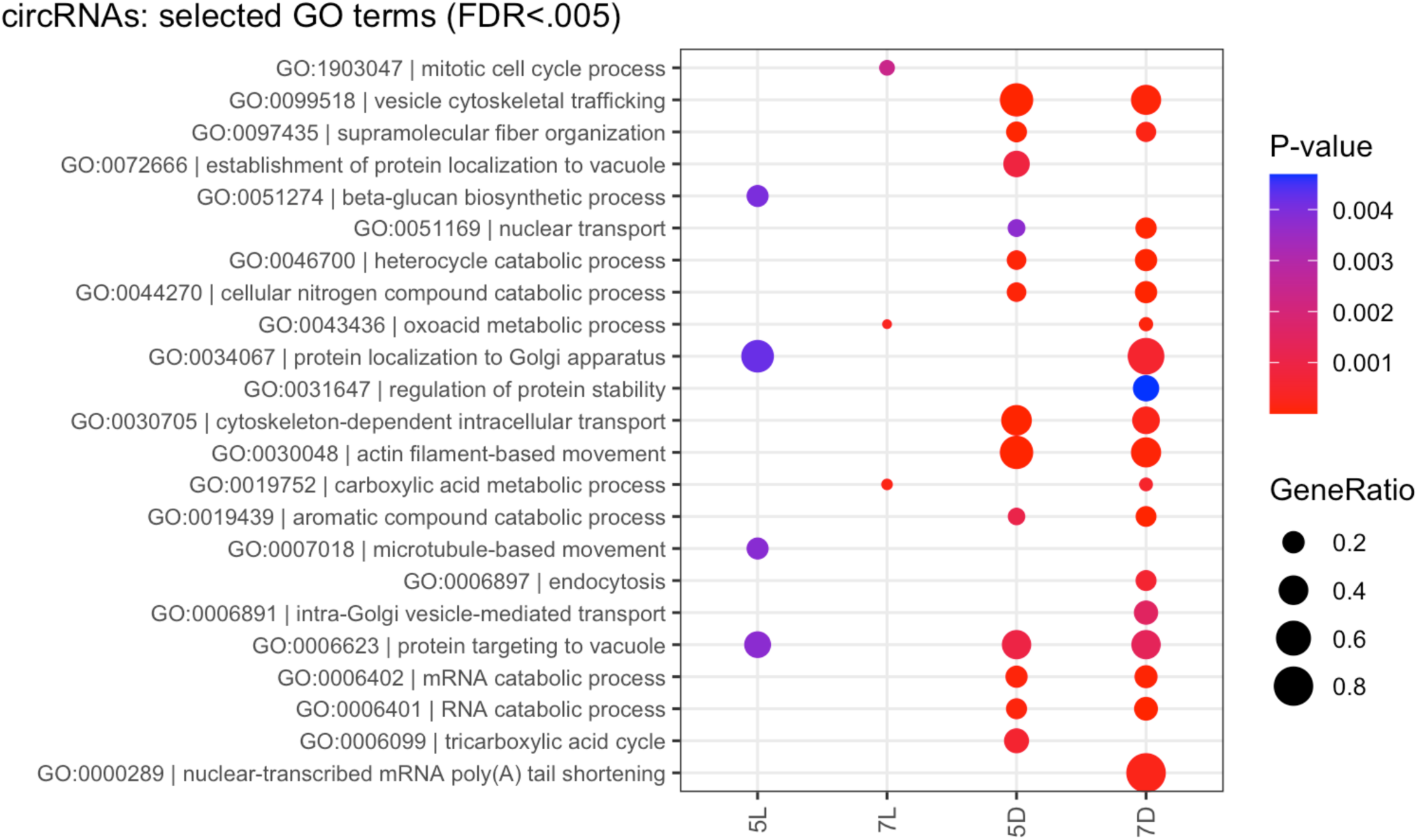
GOterm enrichment of circRNA gene IDs varies between light and dark treatment. To characterize the potential functional differences in genes with circRNAs under light or dark conditions, we analyzed the GOterm enrichment for biological functions of the respective Arabidopsis homologs of the camelina genes. Select GOterm categories with significant (FDR<0.005) enrichment (> 2-fold) that showed fold differences between the light and dark treatment. The gene ratio represents the number of circRNA generating cognate genes in that GOterms divided by the number of total genes in that GOterm. The full PANTHER output is in Supplemental Table S6.

Based on the functional annotations of their cognate genes, most circRNAs (as classified by their cognate gene IDs) showed the expected representation of functional categories (i.e. photosynthesis, amino acid metabolism, etc.) when accounting for the total number of genes represented in our transcriptome. However, several categories that were treatment-specific were also overrepresented. Significant (FDR<0.005) over-representation of GOterms were mostly found in the two dark-treatment groups. CircRNAs for “actin-filament based movement” (GO:0030048) were found for 9 different Myosin homologs, including one Kinesin-like protein, many of which were also included in the overrepresentated GOterm “vesicle cytoskeletal trafficking”, whereas “protein targeting to vacuole” included different vacuolar sorting-associated proteins, such as Transducin, and a vacuole sorting receptor among others, and were also overrepresented but in both dark treatments as well as the 5L treatment. The transcripts contributing to the overrepresentation of the “tricarboxylic acid cycle” (GO:0006099) included multiple isoforms (mitochondrial and peroxisomal) citrate synthase, Aconitases, Isocitrate dehydrogenase and the succinyltransfer subunit of the 2-oxoglutarate DH complex. All of these enzymes would be important for the involvement of seed oil breakdown and conversion to energy in etiolated seedlings.

RNA (and mRNA) catabolic process genes that were overrepresented in the category included different genes for Cap-binding, Decapping enzymes and associated proteins, Ribonucleases, as well as two subunits of the transcriptional regulatory complex CCR4-NOT. Other interesting GOterms with overrepresentation only in the 5L treatment include the “beta-glucan biosynthetic process” (GO:0051274) which included cellulose synthase isoforms (CESA1, CESA3, CESA6) and cellulose synthase-like proteins (CSLE1, CSLB3), and the category “protein localization to Golgi apparatus” (GO:003067) which included Pleckstrin homology (PH) domain-containing protein and two vacuolar protein sorting-associated proteins.

CircRNA from cognate genes that were specifically overrepresented at both 7-day treatments only (7L, 7D) and included in the GOterm for “carboxylic acid metabolic process” (GO:0019752) and “oxoacid metabolic process” (GO:0043436) included 19 genes expressing enzymes of the fatty acid oxidation pathway (Acyl-CoA Oxidase, Isocitrate DH, Aconitase), amino acid biosynthesis (Alanine aminotransferase, Glutamate Synthase (GLS), Glutamine Synthetase (GLN), Arginase) as well as PEP-carboxykinase, Trehalose phosphate synthase, and the plastidal isoforms of Pyruvate kinase and Fructose bisphosphate aldolase 2. Many of these are key enzymes of primary carbon and nitrogen metabolism that generate circRNA in those 7-day old seedlings.

### 3.8. Lipid metabolism genes with circRNA in Camelina

Camelina’s primary use in agriculture as an oil seed crop is for the production of transportation fuel from fatty acids in oil. Although the relevant tissues for the analysis of circRNA involvement in lipid metabolism would obviously be in developing seeds, etiolated and de-etiolated seedlings contain both biosynthesis and degradation pathways for fatty acids and lipids. We used the Arabidopsis lipid metabolism database (Beisson et al. 2003; https://www.arabidopsis.org/browse/genefamily/acyl_lipid.jsp) to identify orthologs in camelina for lipid and fatty acid metabolism. From our data and the publicly available circRNA data from Arabidopsis and other plants circRNA databases, we identified circRNAs associated with fatty acid and lipid metabolism in camelina seedlings. We used those to further compare circRNA resources from rice, and soybean (Table 6, Supplementary Table S7). In camelina we found circRNA for genes of several key metabolic enzymes in the prokaryotic and eukaryotic fatty acid and lipid metabolism pathways. These include intronic as well as exonic circRNAs.

**TABLE 5:**
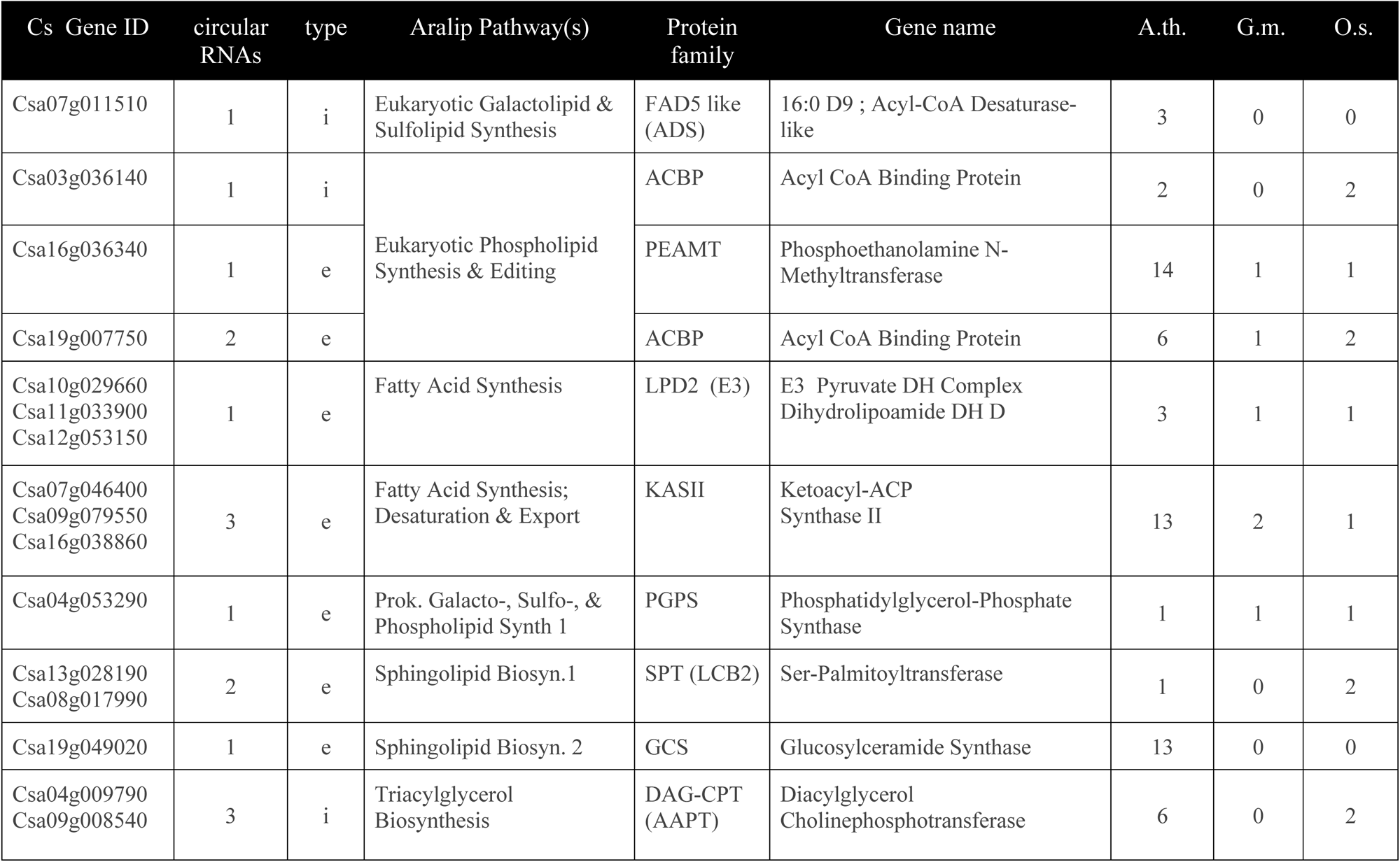
Select lipid and fatty acid metabolism genes with circRNA from Camelina sativa, Arabidopsis, Oryza sativa (japonica), and Glycine max. Data from Arabidopsis, O. sativa, and G. max were identified from PlantcircBASE. CircRNA type refers to intronic (i) or exonic (e). A complete table with orthologs and circRNA BSJ genomic coordinates is available as Supplementary Table S7.

| Cs Gene ID | circular RNAs | type | Aralip Pathway(s) | Protein family | Gene name | A.th. | G.m. | O.s. |
| --- | --- | --- | --- | --- | --- | --- | --- | --- |
| Csa07g011510 | 1 | i | Eukaryotic Galactolipid & Sulfolipid Synthesis | FAD5 like (ADS) | 16:0 D9 ; Acyl-CoA Desaturase-like | 3 | 0 | 0 |
| Csa03g036140 | 1 | i | Eukaryotic Phospholipid Synthesis & Editing | ACBP | Acyl CoA Binding Protein | 2 | 0 | 2 |
| Csa16g036340 | 1 | e |  | PEAMT | Phosphoethanolamine N-Methyltransferase | 14 | 1 | 1 |
| Csa19g007750 | 2 | e |  | ACBP | Acyl CoA Binding Protein | 6 | 1 | 2 |
| Csa10g029660<br>Csa11g033900<br>Csa12g053150 | 1 | e | Fatty Acid Synthesis | LPD2 (E3) | E3 Pyruvate DH Complex Dihydrolipoamide DH D | 3 | 1 | 1 |
| Csa07g046400<br>Csa09g079550<br>Csa16g038860 | 3 | e | Fatty Acid Synthesis; Desaturation & Export | KASII | Ketoacyl-ACP Synthase II | 13 | 2 | 1 |
| Csa04g053290 | 1 | e | Prok. Galacto-, Sulfo-, & Phospholipid Synth 1 | PGPS | Phosphatidylglycerol-Phosphate Synthase | 1 | 1 | 1 |
| Csa13g028190<br>Csa08g017990 | 2 | e | Sphingolipid Biosyn.1 | SPT (LCB2) | Ser-Palmitoyltransferase | 1 | 0 | 2 |
| Csa19g049020 | 1 | e | Sphingolipid Biosyn. 2 | GCS | Glucosylceramide Synthase | 13 | 0 | 0 |
| Csa04g009790<br>Csa09g008540 | 3 | i | Triacylglycerol Biosynthesis | DAG-CPT (AAPT) | Diacylglycerol Cholinephosphotransferase | 6 | 0 | 2 |

### 3.9. Confirmation of circRNAs from lipid metabolism homeologs of KAS II

Circular RNA identification based on short-read sequencing relies on reads that contain identifiable sequences from both, the downstream 5’ splice site covalently linked to an upstream 3’ splice site generating the BSJ of a circRNA. Several events can lead to a “false positive” BSJ read, including genomic rearrangement, trans-splicing, or template switching of the reverse transcriptase (Dodbele et al., 2021; Nielsen et al., 2022). Specific circRNAs can be validated with RT-PCR using divergent and convergent primer sets on cDNA and genomic DNA (gDNA) templates from the respective plant tissue (Figure 11A). After separating the products by gel electrophoresis (Figure 11B), we confirmed the sequences of the amplification products by Sanger sequencing. Divergent primers should only be able to amplify circRNA from the cDNA and produce no product from gDNA amplification. Amplification of a product with the same BSJ from gDNA indicates genomic duplication or rearrangements.

**Figure 11:**
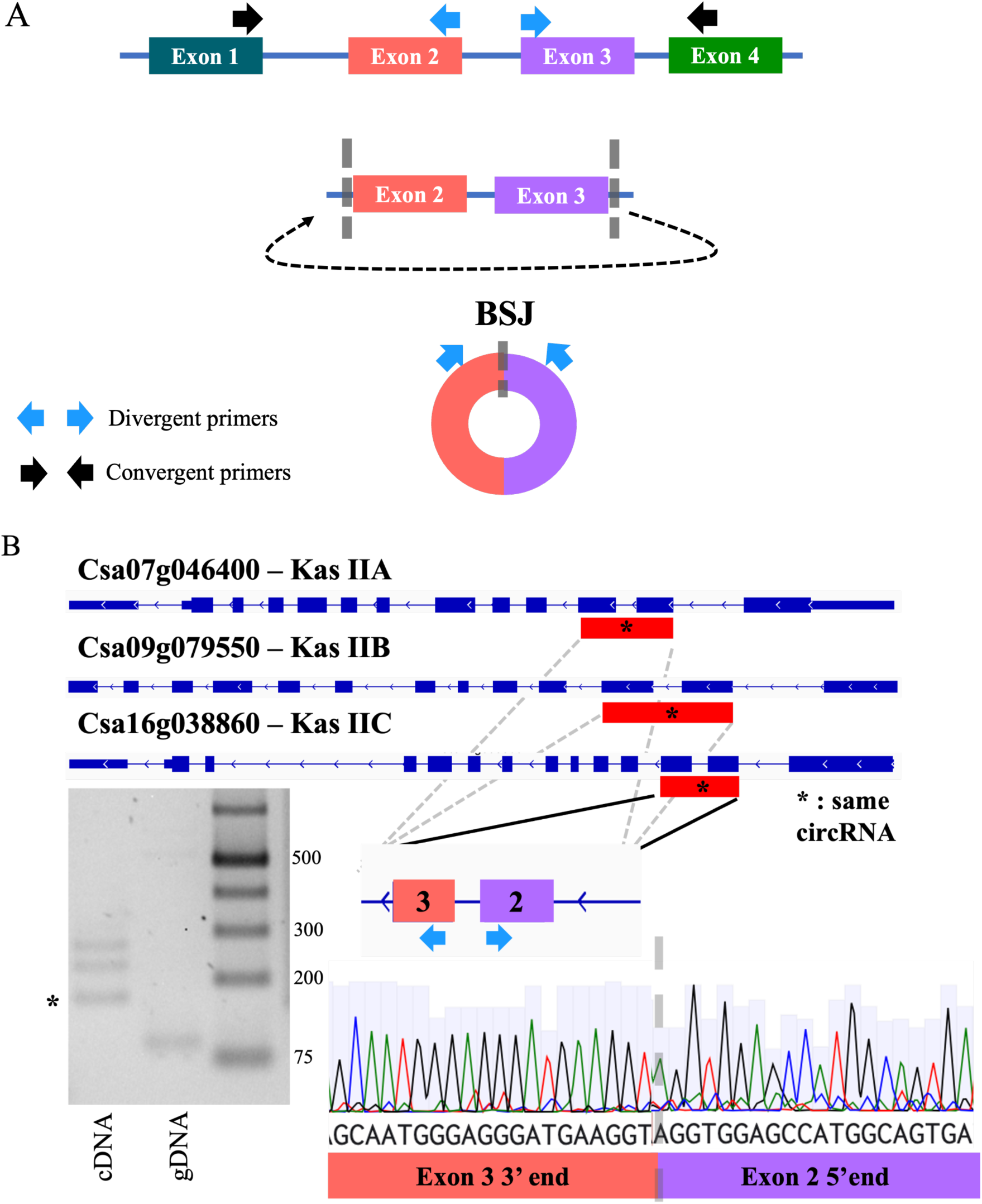
Confirmation of circRNA from Kas IIA, Kas IIB, and Kas IIC. CircRNA was confirmed for Kas IIA, Kas IIB, and Kas IIC by using divergent and convergent primer pairs (A) to generate cDNA for PCR amplification and sequencing. (B) gDNA was used as a control. A full list of primers tested and full image of gel electrophoresis in Table S1 and Figure S1.

KASII has three homeologs, one in each of the three subgenomes: Csa07g046400 (KASII A), Csa09g079550 (KASII B), and Csa16g038860 (KASII C). Two putative circRNAs originated from these genes. We validated one circRNA which contains exons 2 and 3 of KASII A/B/C with corresponding BSJ coordinates Chr7:23795661-23796179, Chr9:30060624-30061144, Chr16:19858862-19859379. These KASII homeologs have very similar sequences over the circRNA region and we were unable to differentiate between them in PCR and Sanger sequencing results. Thus, it is unclear at present which or how many of these homeologs express the circRNA. A second putative circRNA was identified from KASII B but sequence quality was not sufficient to conclusively validate it, potentially due to mixed populations of amplicons from the different subgenomes. KASII also has orthologs in the legume *Lotus japonicus* and two close orthologs, LotjaGi4g1v0321700 and LotjaGi6g1v0221600 also express circular RNAs (Franklin, Utley, Budnick - manuscript in preparation).

### 3.10. CircRNA with putative miRNA binding sides

CircRNAs have been shown to bind miRNA and can serve as competing endogenous RNAs to reduce degradation of the linear transcripts that are miRNA targets (Zhou et al., 2021). In Camelina, miRNAs have been identified and some of their targets have been predicted based on sequence analysis (Poudel et al., 2015). The functional involvement of miRNA in lipid metabolism has also been experimentally confirmed, showing that overexpression of miRNA167 in camelina results in changes in the lipid profile and seed size (Na et al., 2019). We compared the previously sequenced camelina miRNAs from Poudel et al. (2015) (Poudel et al., 2015) and their predicted targets to the circRNAs identified in our study here (Supplementary Table S8).

Eighty-seven of the genes predicted to be targeted by miRNA in Poudel et al. (2015) were found to express circRNAs in our data. Genomic sequence was extracted between the back splice junction coordinates of the 97 unique circRNAs corresponding to the 87 unique genes and miRNA targets were predicted using psRNATargetV2 (Dai et al., 2018). This revealed 101 putative miRNA target regions across 45 circRNA genomic sequences.

One trend in this target prediction set is a high number of genes implicated in cytoskeletal features. Among these were six circRNAs from putative P-loop containing nucleoside triphosphate hydrolase genes, 3 circRNAs from 2 HEAT repeat containing genes, and two circRNAs from a single putative myosin 2 gene; annotations are based on orthology to arabidopsis. The miRNAs for these genes showed some overlap which could imply a regulatory network connecting circRNAs and miRNAs in the camelina cytoskeletal system (Figure 12).

**Figure 12:**
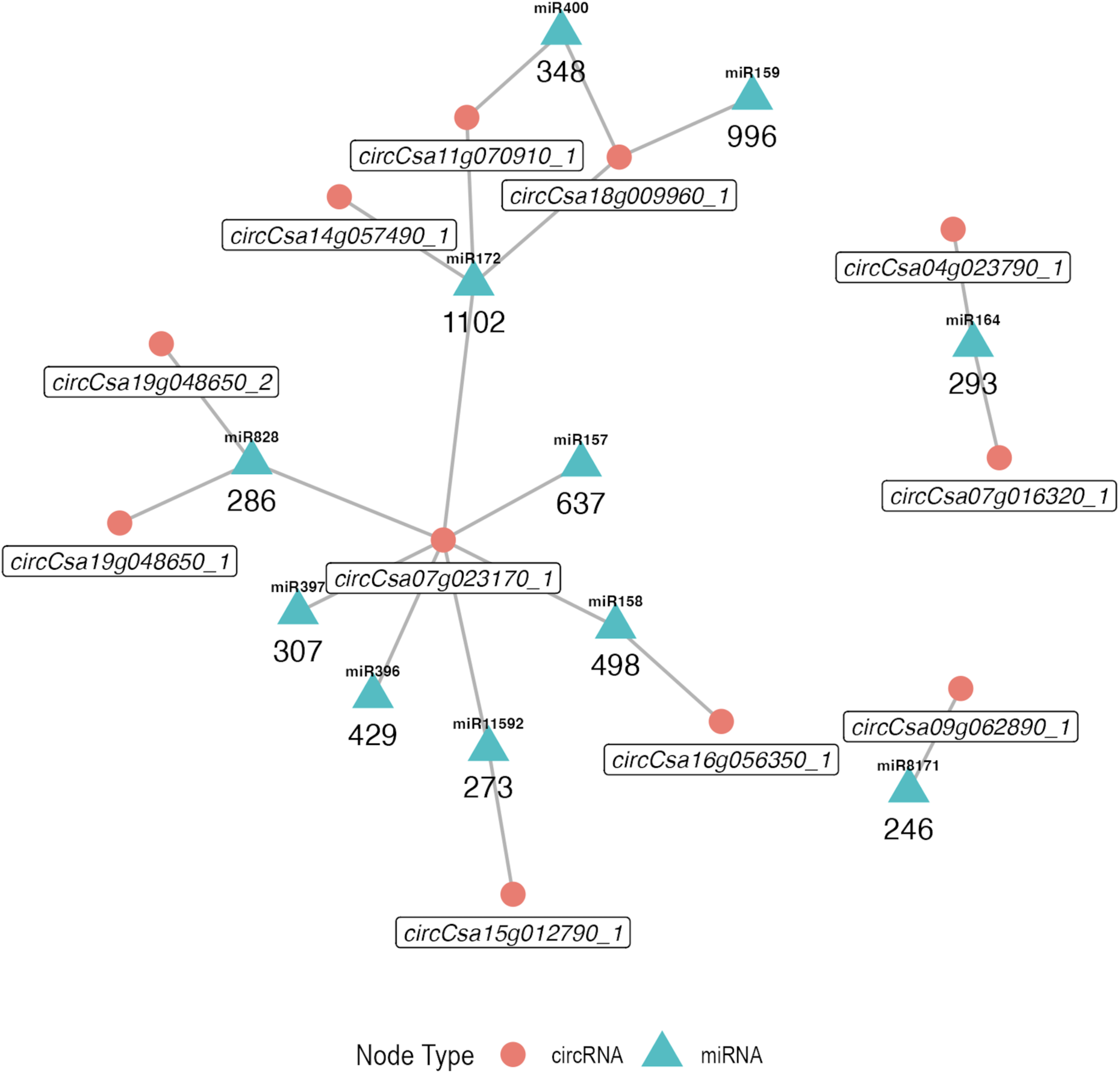
Network visualization of miRNA:circRNA connections from genes associated with the cytoskeleton. A network was constructed based off of circRNAs which are putative targets for *C. sativa* miRNAs. Circular RNA nodes are displayed as red circles and miRNA nodes are displayed as blue triangles. The number of predicted mRNA targets is shown below each miRNA.

This network of predicted miRNA targeting implicates circCsa07g023170_1 from Csa07g023170, which shows orthology to the arabidopsis gene ILA (AT1G64790), as a potential hub with predicted targets for 7 distinct miRNAs, 4 of which are predicted to target other circRNAs expressed in this set of cytoskeletal related genes.

### 3.11. Incorporating circRNA tracks in CamRegBase

To make the Camelina circRNA readily available to the community, we incorporated it into the CamRegBase (https://camregbase.org/) knowledge base (Gomez-Cano et al., 2020). The circRNA data for each environmental condition was converted into GFF Generic Feature Format (G. Pertea & Pertea, 2020; Stein, L., 2013). The GFF files are hosted on the database as static FTP directories using standard Apache web hosting software (Fielding & Kaiser, 1997). CircRNA data in GFF format is made accessible to users through the JBrowse user interface (Buels et al., 2016). In JBrowse, circRNA reads can be visually aligned with other searchable CamRegBase annotations and data tracks. Throughout the CamRegBase website, hyperlinks take users directly to regions of interest in their customized JBrowse interface.

The JBrowse web-based GUI allows users to broadly search and explore the genome, zoom into individual base pairs and customize display settings, export circRNA data, or add their own local data to the display (Buels et al., 2016). The visualization, customization, and all interactive features are powered by local javascript and cookies on the user’s computer, simplifying the server hardware requirements. Perl scripts (Wall, 1994) were used to encode text indices in the form of directory structures to support intelligent search/autocomplete of the JBrowse search bar using only static file hosting and again no server-side control was necessary.

**Figure 13:**
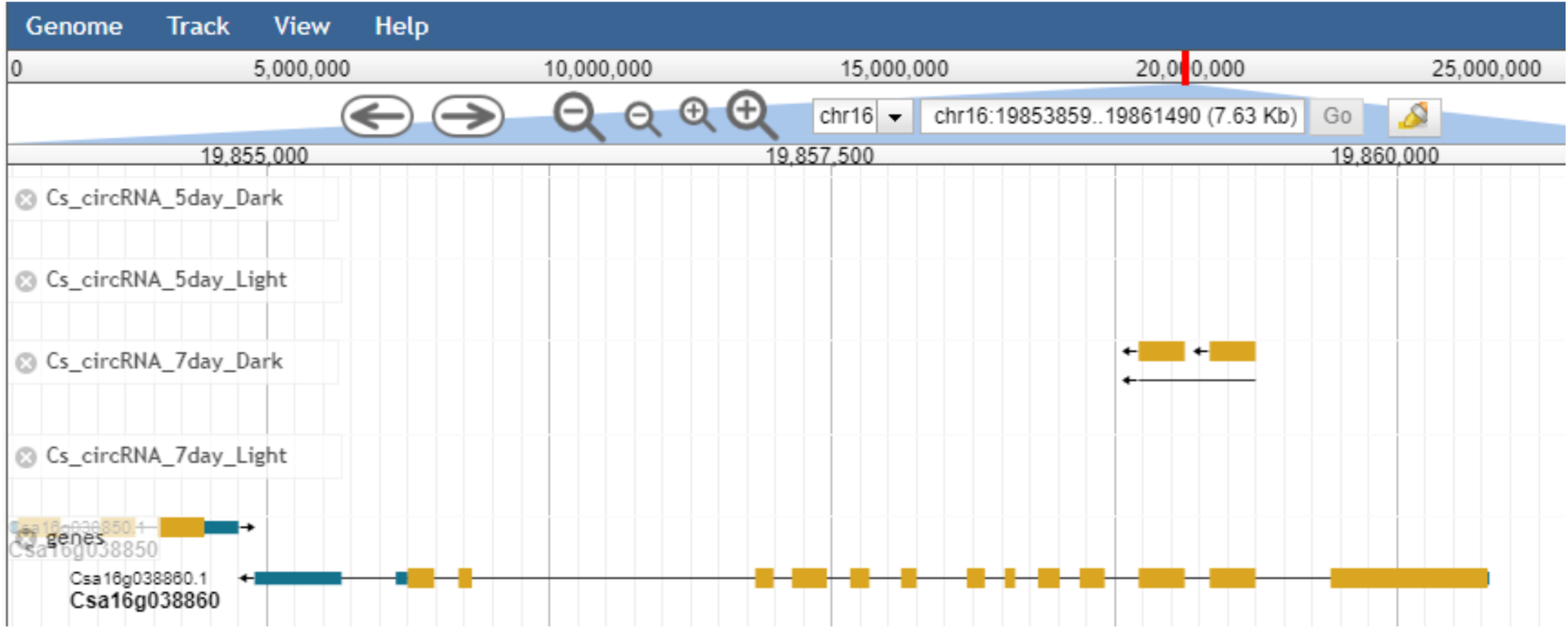
Screenshot of CircRNA tracks identified in the 4 treatments and alignment to KAS IIC Csa16g038860.

## 4. DISCUSSION

Regulation of gene expression occurs on many levels. It has become clear from studies in animal and plants that circRNA is involved in this regulation at both, the transcriptional and post-transcriptional level. We identified 3,447 genic and 307 intergenic putative circRNAs from etiolated and de-etiolated *Camelina sativa* seedlings. While this study provides a first landscape of circRNA in camelina seedlings, verification of these circRNAs and identification of circRNAs formed under other abiotic and biotic conditions and in different tissues during development will be necessary. Few high-throughput methods currently exist to evaluate the different functions these circRNAs on transcriptional and posttranscriptional regulation of gene expression, especially with respect to lipid and oil metabolism in camelina seeds. We discuss below the challenges of circRNA detection and validation as well as their potential functions in plant development and lipid metabolism.

### 4.1. Challenges to circRNA identification

The identification of true circRNAs using high-throughput sequencing technology is still a challenge (Drula et al., 2024). The low abundance of circRNA of less than 0.1% of non-ribosomal RNA requires deep sequencing. We have shown here that the sequencing depth (>60 million reads/sample) was not limiting in the detection of unique circRNAs in rRNA-depleted cDNA libraries, generated with random primers. Other methods that enrich circRNA prior to sequencing by degradation of linear RNA with RNAse R or polyA-pull down treatment can yield higher numbers of unique circRNAs, but different pools of circRNAs are identified with short read sequencing (Philips et al., 2020) compared to long-read sequencing using nanopore technology (Budnick et al., 2024). It is not clear whether these differences in circRNA identification by different methods is due to specific molecular features that those different populations of circRNA have such as their locations/accessibility, secondary structure, interactions with other molecules, or overall stability. Because the use of random hexamer primers is necessary to identify circRNA (due to its lack of a polyA tail), a GC bias in random hexamer primering is possible, even likely (K. D. Hansen et al., 2010). Another known concern for amplification of false splice junction reads is the ability of reverse transcriptases to switch templates during cDNA synthesis. The in-vitro template switching mechanism during cDNA synthesis can be reduced by degradation of linear RNA prior to RT and/or by using optimized reverse transcriptases that have reduced template-switching ability (Cocquet et al., 2006; Dodbele et al., 2021).

Another challenge is the computational detection of BSJ in large populations of sequence reads generated by random primer reverse transcription. CircRNAs are detected from sequencing reads by selecting reads that do not align to the genome and check for discontiguous alignments and changes in orientarion. Different bioinformatics pipelines have been developed to optimize the precision and sensitivity of circRNA detection. A recent benchmarking study of circRNA detection programs showed that the precision of the detected circRNA was very high (over 95%), but the different tools varied almost 50-fold in the sensitivity of detection, which results in the actual number of circRNAs they identify as such in a given sample (Vromman et al., 2023). CIRI and CLEAR pipelines used in our study (Gao et al., 2015; Ma et al., 2019) showed high precision and fairly high sensitivity, although CIRI has a read cut-off of at least 2 reads per circRNA which CLEAR does not have. We have confirmed true circRNA from several predictions in CLEAR with only a single read count, including KAS II (Figure 11; unpublished data). No single tool identified all true circRNAs (Zeng et al., 2017), which is why it is standard procedure to use at least two bioinformatic algorithms to detect circRNAs and reduce the number of false positives, a practice which was performed in this study (T. B. Hansen, 2018). However tedious, this standard has led to new tools that have emerged which combine multiple circRNA detection methods (Gaffo et al., 2022) in order to identify most-likely candidates for circRNA validation. In our investigations we also found that several false positive BSJ were caused by actual unannotated sequence repeats in the DNA that were not present in the published reference genome database.

In animal systems, several inverse repeat sequences flanking circRNAs have been shown to be essential in their biogenesis (Jeck et al., 2013; D. Liang & Wilusz, 2014) and are therefore possible sequence markers that would allow for the sequence-based identification of putative RNAs. Far less is understood in plant circRNA biogenesis, but recently miniature inverted-repeat transposable elements (MITE) insertions into flanking sequences have been identified in a few circRNA flanking regions in maize and poplar (Han et al., 2020; Song et al., 2021).

### 4.2. The many functions of circRNA

Mechanisms of circRNA biogenesis, including their inducibility and variety of different functions in gene expression are still poorly understood. In plants, circRNA function has been demonstrated in alternative splicing (Conn et al., 2017), miRNA sequestration (J. Zhang et al., 2020), and antisense suppression (H. Zhang et al., 2021).

Alternative splicing is a widespread mechanism in eukaryotic organisms that expands the repertoire for proteins and their functions during development and in response to biotic or abiotic stress. While many versions of transcripts from a single gene are caused by alternative start and stop codons, alternative splicing is an inducible, co-transcriptional process that requires the activity of the spliceosome. In Arabidopsis, about 80% of nuclear-encoded genes contain introns and under “normal” growth conditions, about 61% of the multi-exonic transcripts are alternatively spliced (Marquez et al., 2012). Stress-specific regulation of alternative splicing has been shown in Arabidopsis for pathogen defense (Howard et al., 2013), Fe-deficiency and phosphate stress (W. Li et al., 2013), and tissue specificity (Martín et al., 2021; Misra et al., 2023) with the majority (40%-60%) of alternative splicing events across conditions representing intron retention. A comparison of alternative splicing events across genomes between *Arabidopsis thaliana* and *Camelina sativa* showed about 70% conservation for the four different AS categories of exon skipping, intron retention and alternative splice donor or acceptor sites (Martín et al., 2021).

The only conclusive evidence showing circRNA as a cause of alternative splicing is the exon skipping event in the Arabidopsis gene *Sepallata3* that causes a change in the number of petals (Conn et al., 2017). The authors showed that a circRNA derived from exon 6 of the Sepallata3 gene can increase the abundance of the cognate exon-skipped alternative splice variant, resulting in a flower petal phenotype. The overexpression of circRNA, but not linear RNA from exon 6 resulted in an increased petal count and decreased stamen count. SEP3 exon 6 circRNA formed an R-loop with its cognate gene DNA, detected by the R-loop specific S9.6 antibody (Conn et al., 2017). Genome-wide analysis of R-loop formation with circRNA can be captured by DNA:RNA immunoprecipitation with the S9.6 antibody followed by high-throughput sequencing (DRIP-seq). A recent study using DRIP-seq in Arabidopsis showed that the majority of genes formed R-loops (86%). R-loops were formed with both, sense and antisense strand DNA and relatively short, mostly between 100-500 bp long. Both sense and antisense R-loop formation was correlated with transcription and associated with various histone modifications. The wide-spread formation of R-loops at GpC islands is thought to protect against de-novo DNA methylation to regulate transcription (Xu et al., 2017).

Genome-wide identification of circRNAs in R-loop formation in poplar differentiating xylem was identified by DRIP following the digestion of linear RNA by RNAse R, an exonuclease to which circRNA is resistant. The study identified 181 circRNAs (4.4% of total circRNAs) and 672 AS events that overlapped with R-loops in this tissue and confirmed the causative effect of AS through overexpression of circRNA for select genes, including a hemicellulose synthase isoform (X. Liu et al., 2021). Interestingly, the formation of R-loops to affect expression and alternative splicing would theoretically only require two molecules of a specific circRNA in a diploid cell, which could explain the very low abundance of some circRNAs seen in our data as well as other plant circRNA sequencing studies.

So far, few miRNA-binding circRNAs have been experimentally verified in plants. One study found an intronic rice circRNA (*Os06circ02797*) involved in seedling growth that can bind *OsMIR408*. This was evidenced by RT-qPCR of CRISPR-Cas9 deletion mutants showing an increase of *OsMIR408* and downregulation of *OsMIR408*-targets compared to wild type (Zhou et al., 2021). Another study in bamboo showed a miRNA-circRNA-mRNA network, responsive to nitrogen fluctuations. This was supported by a dual-luciferase reporter assay that showed the miRNA bound to a synthesized version of the circRNA (Zhu et al., 2023). In *Arabidopsis*, a competing endogenous circRNA that sequesters a miRNA away from a drought tolerance gene leads to drought resistance via functional circRNA overexpression and localization tests (Yin et al., n.d.). Other circRNAs have been predicted to be ceRNAs, including circRNAs involved in defense response in tomato to *P. infestans* (Hong et al., 2020). We identified a possible miR472 target site in Rubisco small SU (Csa12g084210) as well as its circRNA. The Rubisco small SU circRNA as well as its cognate linear RNA was more highly expressed in the light samples compared to the dark samples (Figure 8D). Interestingly, the other two RubisCO small SU homeologs did not generate circRNAs. However, we did not sequence miRNAs from these samples and therefore a putative miRNA:circRNA interaction is speculative.

As with many high-throughput omics approaches on whole seedlings or tissues, a big unknown is the cellular and subcellular (co-)localization of effectors, regulators, and targets. It has been shown that miRNAs, linear RNAs, proteins, peptides, and viral circular RNA are mobile throughout the plant - between cells, tissues, and organs. If circRNA is also a mobile messenger molecule has not yet been established. CircRNAs were identified in apoplastic fluid and are thought to be involved in defense and signaling (Zand Karimi et al., 2022).

### 4.3. CircRNA in lipid metabolism

Many of the genes involved in fatty acid and lipid biosynthesis have been shown to generate circRNA. In Arabidopsis, most of the key pathway genes in the prokaryotic and eukaryotic lipid biosynthesis pathway as well as in lipid and fatty acid degradation have BSJ fragments identified in the circRNA database (http://www.deepbiology.cn/circRNA/). While economically relevant oil biosynthesis and accumulation occurs in the seeds, most enzymes are active in other tissues to generate membrane lipids or precursors for other molecular groups including hormones and waxes (Li-Beisson et al., 2013). Potential networks of circRNA:miRNA as competing endogenous RNA (ceRNA) networks have been identified in soybean fatty acid synthesis (Ma et al., 2019).

De novo synthesis of fatty acids occurs in the plastid (Li-Beisson et al., 2013). The major substrate, acetyl-CoA is synthesized by the Pyruvate Dehydrogenase Complex (PDHC) from pyruvate. The PDHC contains three components: E1 (pyruvate dehydrogenase), E2 (dihydrolipoyl acyltransferase), and E3 (dihydrolipoamide dehydrogenase, LPD). We identified putative circRNA from the E3 component of PDHC (Table 6). The next step in the de novo fatty acid biosynthesis is the condensation between two acetyl-CoA molecules to malonyl-CoA by the Acetyl-CoA Carboxylase (ACCase). We identified BSJ reads for a putative circRNA for ACCase (Supplementary Table S3) but attempts to confirm the circRNA of ACCase provided inconclusive results. We therefore are not listing ACCase in Table 6, although circRNAs were found for ACCase in Arabidopsis. Malonyl-CoA is transferred to Acetyl-Binding Protein (ACP) for further rounds of elongation of the carbohydrate chain with Acetyl-ACPs by the Fatty Acid Synthase complex. The first enzyme in that complex is KetoAcyl-ACP Synthase III (KASIII) which catalyzes the condensations of Malonyl-ACP with Acetyl-CoA to increase the length of the hydrocarbon chain by 2-carbon units. Further rounds of acyl-ACP condensation with acetyl-CoA are catalyzed by KASI until a C16:0-ACP chain is formed. The final condensation reaction extending C16:0 to C18:0 is catalyzed by KASII. We clearly identified circRNA spanning exons 2 and 3 of KAS II. The function of this circRNA is not determined and we were not able to identify any miRNA binding site. There are also no reports on alternative splice versions of KASII in camelina or Arabidopsis that encompass exons 2-3. However, we identified multiple alternative transcripts for KAS IIB and KAS IIC that show exon-skipping, intron-retention and also an antisense fragment identified by the splice-aware aligner used in this study (Supplementary Figure S2; Supplementary Table S5).

Termination of the fatty acid extension is catalyzed by transfer from ACP to CoA, catalyzed by acyl length specific fatty acid thioesterases (FAT) enzymes. As a bioenergy oil crop for drop-in replacement fuel, shorter and saturated fatty acids are more desirable to reduce processing energy and reduce oxidation. Several engineering approaches targeting thioesterases and KASII were successful in modifying the fatty acid profile of camelina seed oil to include shorter, more saturated fatty acids (Hu et al., 2017; H. J. Kim et al., 2015; Pidkowich et al., 2007). Engineering of circRNAs could provide another mechanism to achieve this goal. Ultimately, circRNAs from the relevant tissues during seed development need to be sequenced for circRNA to characterize their functions during seed development and oil accumulation.

## 5. CONCLUSION

CircRNA is an interesting new class of regulators of gene expression in eukaryotes. Their implication in alternative splicing, transcript stability and translational efficiency in plants calls for more investigations into the mechanisms of biogenesis and control of the inducibility of circRNA formation. The use of circRNAs in plant protection or engineering applications will require more knowledge about their functions, however, the use of RNA in crop production is an interesting new area of commercial interest. CircRNAs could also serve as biomarkers for abiotic or biotic stress in plants. In human cell lines circRNAs have been identified that are specific enough to act as a diagnostic biomarker in the detection of specific cancers like non-small-cell lung cancer [75]. First tests in mouse models for muscle atrophy injecting circRNAthat acts as a miRNA sponge have shown improvement of pathological symptoms (J. Li et al., 2022). Other applications using engineered endogenous or exogenous circRNA are being developed and the first circRNA vaccines are being tested (Qu et al., 2022). This first identification of circRNA from *Camelina sativa* provides a basis to investigate the functions and potential for engineering of lipid metabolism using circRNA as a new player in the regulation of gene expression.

## 6. SUPPLEMENTARY MATERIAL

**Supplementary Figure S1:** Full gel electrophoresis image of KASII circRNA validation corresponding to Figure 11.

**Supplementary Figure S2:** Abundances of transcripts and alternative splice variants identified from Supplementary Table S5 for the three KAS II homeologs.

**Supplementary Table S1:** RT-PCR primer sequences used for validating KASII circRNA. Primers are labeled with lane numbers corresponding to the original gel electrophoresis images in Supplemental Figure S1.

**Supplementary Table S2:** Sample metadata. Includes the number of read pairs (Q>30) for each sample.

**Supplementary Table S3:** CIRI2 and CLEAR pipeline outputs and comparisons. This supplemental table contains all CLEAR (Sheet 1) and CIRI2 (Sheet 2) outputs, as well as the clustered and merged circRNA used in final analysis (Sheet 3).

**Supplementary Table S4:** Normalized counts for linear transcripts. This table also contains calculations used in determining DEGs.

**Supplementary Table S5:** Initial output of Ballgown statistical analysis package. Sheet 1 description of symbol designation of “t.class” column in original output of Ballgown (Sheet 2).

**Supplementary Table S6:** Full list of GOterms corresponding to Figure 10.

**Supplementary Table S7:** Complete table from Figure 6, comparing orthologs and circRNA BSJ genomic coordinates from arabidopsis, rice, and soybean.

**Supplementary Table S8:** Combined table of miRNAs and circRNAs for the subset of genes found to have targeting miRNAs by Poudel et al. which also contained circRNAs in our data.

## 7. DATA AVAILABILITY

All sequencing data and corresponding sample information has been deposited on the Dryad server for free public access: Effect of light exposure on circular RNA and alternative splicing in Camelina sativa https://doi.org/doi:10.5061/dryad.m0cfxpp8k. Custom R scripts used for the formatting and analysis of data are available via Github: https://github.com/SederoffLab/Camelina_circRNA. Other data are available in supplementary materials or via contacting the authors.

## 8. AUTHOR CONTRIBUTIONS

Brianne Edwards and Delecia Utley contributed equally to this study. **Delecia Utely:** Data curation; formal analysis; methodology; software; validation; visualization; writing. **Brianne Edwards:** experimental design, investigation, Data curation; formal analysis; methodology; software; review and editing. **Asa Budnick**: Data curation; formal analysis; software methodology; visualization; writing—review and editing. **Erich Grotewold**: methodology; validation; writing—review and editing. **Heike Sederoff:** Conceptualization; formal analysis; funding acquisition; investigation; project administration; supervision; visualization; writing— original draft; writing—review and editing.

## 10. ACKNOWLEDGEMENTS

We thank Oliver Tessmer for incorporating circRNA loci information into CamRegBase. Funding for this work was provided by the Department of Energy, Office of Science (DE-SC0018269 to H.S. and E.G. and DE-SC0022987 to E.G.), the Novo Nordisk Foundation (InRoot NNF19SA0059362), a National Science Foundation NRT Fellowship (DGE-1828820 to A.B., and D.U.), and a Southern Regional Educational Board Fellowship to D.U.

## Abbreviations

AS: Alternative splice
BSJ: Backsplice Junction
ceRNA: competing endogenous RNA
CircRNA: circular RNA
FP: False positive
FPB: Fragments per Billion mapped bases
RT-PCR: Reverse Transcription Polymerase Chain Reaction
RUBISCO: Ribulose Bisphosphate Carboxylase/Oxygenase

---

**Supplemental Figure S1.**
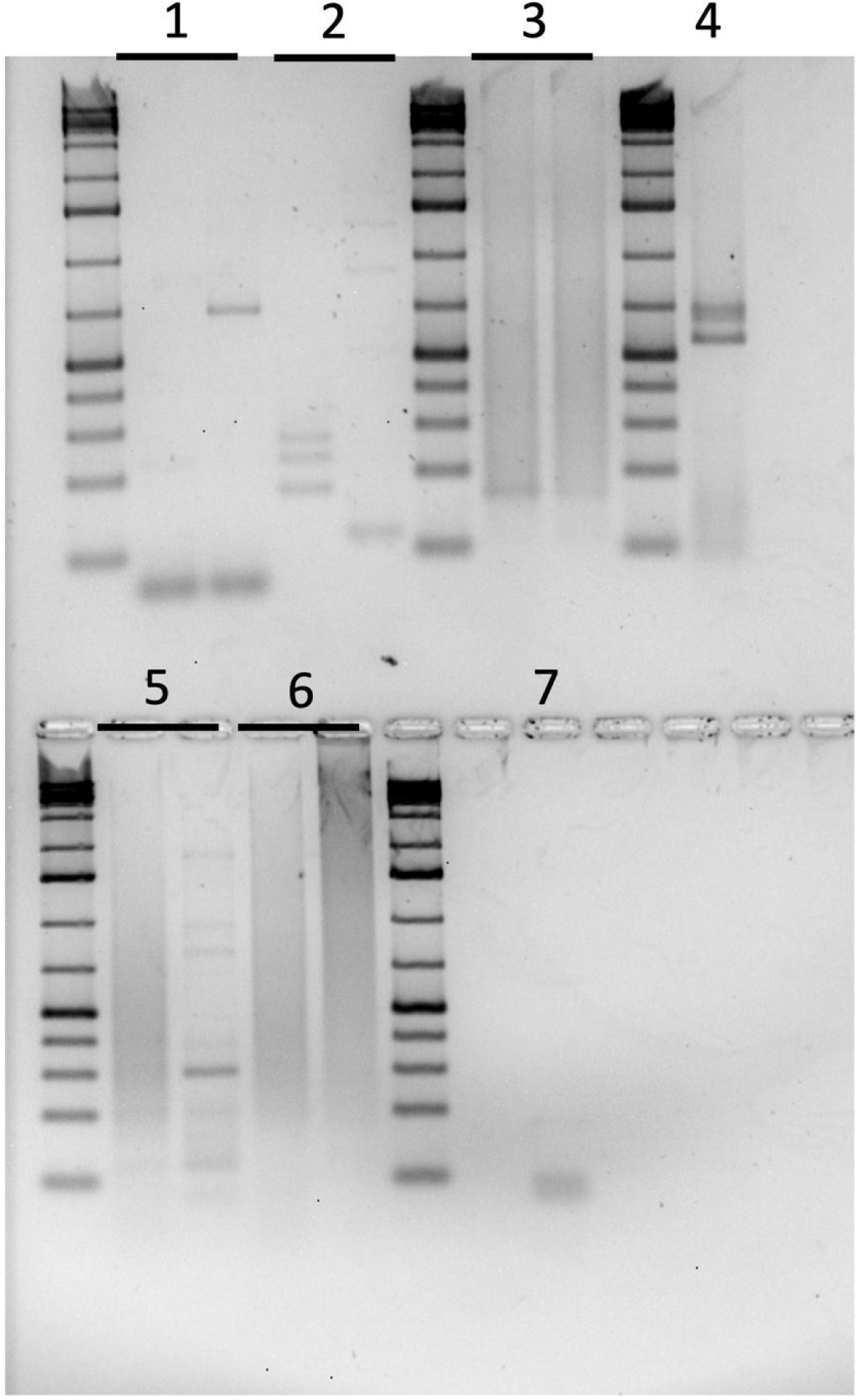
Full gel electrophoresis images of circRNA RT-PCR validations corresponding to Figure 11. Numbers correspond to the RT-PCR/gDNA-PCR reactions. Each underlined number indicates two lanes, the left lane is cDNA template and the right lane is gDNA template (gDNA used as false positive control). Positive linear CsActin2 control and no template controls are shown in lanes 4 and 7, respectively. Note that despite a product being present in no template reaction, the product was extracted for Sanger sequencing and determined to be primer dimer. Gel was prepared as 2% agarose in TAE, run at 130V and post-stained with GelGreen. The ladder used is Generuler 1kb DNA ladder.

**Supplemental Figure S2:**
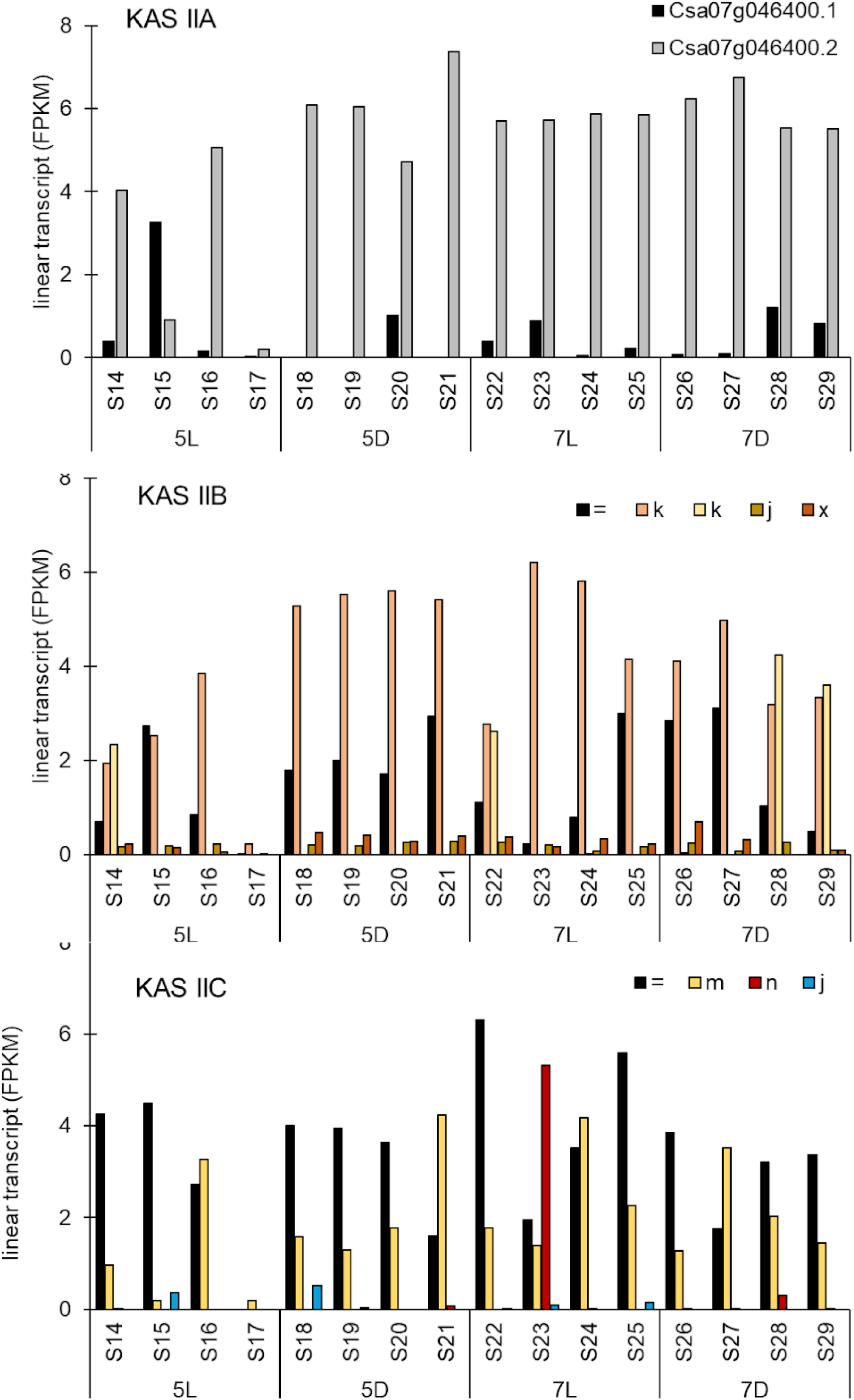
Normalized sequence read numbers for KAS II A-C splice versions. Attributes of novel transcripts were compared to the known, annotated transcripts using GFFcompare (G. Pertea & Pertea, 2020). Differential expression analysis was performed at the transcript level with fragments per kilobase of transcript per million reads sequenced (FPKM) as the expression measurement using the Ballgown statistical analysis package in R (Frazee et al., 2015). While KAS IIA shows higher expression for Csa07g046400.2 than for Csa07g046400.1. Alternative splice versions were also identified for KAS IIB and KAS IIC, with the highest abundance of the complete linear transcript (=) for KAS IIC and a transcript with an additional exon (k) for KAS IIB. None of the alternative splice versions showed treatment-specific abundance changes.

